# Monitoring deep-tissue oxygenation with a millimeter-scale ultrasonic implant

**DOI:** 10.1101/2021.03.05.434129

**Authors:** Soner Sonmezoglu, Jeffrey R. Fineman, Emin Maltepe, Michel M. Maharbiz

**Author notes:** e-mail: S.S.; M.M.M.

## Abstract

Deep tissue oxygenation monitoring has many potential applications. Vascular complications after solid organ transplantation, for example, frequently lead to graft ischemia, dysfunction or loss, and can occur months after transplantation. While imaging approaches can provide intermittent assessments of graft perfusion, they require highly skilled practitioners, and fail to directly assess graft oxygenation. Existing tissue oxygen monitoring systems have many drawbacks, including the need for wired connections, the inability to provide real-time data, and, crucially, an operation that is limited to surface tissues. Here, we present the first wireless, minimally-invasive deep tissue oxygen monitoring system that provides continuous real-time data from centimeter-scale depths in a clinically-relevant large animal (sheep) model and demonstrates operation at great depths (up to 10 cm) through *ex vivo* porcine tissue. The system relies on a millimeter-sized, wireless, battery-free, implantable luminescence oxygen sensor that is powered by ultrasound and capable of bi-directional data transfer with an external transceiver. We present various aspects of system and sensor performance and demonstrate the operation of the system *in vitro* in distilled water, phosphate-buffered saline (PBS) and undiluted human serum, *ex vivo* through porcine tissue, and *in vivo* in a sheep model. We believe this technology represents a new class of diagnostic system particularly suitable for organ monitoring, as well as other surgical or critical care indications.

## Introduction

Tissue oxygenation is a critical determinant of organ function. Our abilities to track it, however, are typically limited to indirect approaches that limit clinical care. Organ transplantation provides a good example. Despite advances in medicine, demand for organ transplantation continues to rise^1^. While highly successful, vascular complications such as hepatic artery thrombosis following orthotopic liver transplantation frequently lead to graft ischemia, dysfunction, and loss^2^. Similarly, renal and pancreatic transplants, along with heart transplants, are frequently complicated by both acute as well as longer-term vascular complications such as graft vascular disease that are the single most important long-term limitation to solid organ transplantation^3^. In the months to years following transplantation, graft arteries characteristically develop severe, diffuse intimal hyperplastic lesions that restrict microvascular flow and cause graft ischemia, ultimately resulting in graft dysfunction. The ability to continuously monitor graft oxygenation status via a wireless sensor placed at the time of transplant would greatly augment our abilities to track tissue perfusion, allowing timely intervention, and help differentiate parenchymal rejection from graft vascular disease in the setting of graft dysfunction.

Most clinically available, direct O_2_ monitoring systems rely on electrochemical sensors (Clark-type O_2_ electrodes)^4, 5^ and luminescent probes^6–8^. Clark-type electrodes consume oxygen, a fact that potentially confounds accurate measurement of tissue O_2_, especially at low O_2_ concentrations, and are not well-suited to long-term measurements due to their susceptibility to biofouling^9^. These facts led to the development of luminescent O_2_ probes as an attractive alternative to Clark electrodes. These probes, although easy to use and accurate, are invasive and fragile, and rely on wired connections, leading to implantation problems and limiting chronic use. Other methods, including fluorine magnetic resonance imaging (^19^F-MRI) spectroscopy^10, 11^ and electron paramagnetic resonant (EPR) spectroscopy^12, 13^, have also been used to directly probe *in vivo* local oxygen content. However, in addition to relying on large, expensive equipment, these methods suffer from an inability to provide real-time O_2_ data as they require long scanning times to quantify O_2_ concentration with sufficient resolution.

As an alternative to the above direct O_2_-sensing approaches, near-infrared spectroscopy (NIRS)^14–17^ has been demonstrated the ability to quantify tissue O_2_ saturation (StO_2_) levels. However, the use of noninvasive NIRS systems for *in vivo* O_2_ detection, especially in deep tissues, is limited by the penetration depth of near-infrared light^14, 15^. NIRS systems with a probe could enable StO_2_ monitoring in targeted regions of deep tissues; however, these systems are invasive and generally rely on fiber-optic tethers^16^ or wired connections^17^, limiting their use in the hospital environment, due in part to increased infection risk. An implantable wireless oximeter that measures StO_2_ has been demonstrated in the literature^18^. That work showed *in vivo* regional StO_2_ measurements from just below the skin in an untethered, freely moving mouse in a home cage; the system relied on near-field electromagnetic (EM) waves for wireless power transfer (restricting it to shallow operation) and the implant occupied ∼100 mm^3^. An implantable system based on an implant of ∼11 mm^3^ has demonstrated electrical stimulation of the heart in an anesthetized rabbit model at 5 cm depth (1 cm air gap and 4 cm-thick tissue); the system relied on a phased array with control circuitry outside of the body which focused EM energy in the mid-field regime to the implant^19^.

Recently, ultrasound (US) power (and data) transfer has been explored as an attractive option for implantable devices deep in tissue^20, 21^, enabling a reduction in their volume to a few mm^3^. In contrast to EM, US allows wireless power transfer and communication via pressure waves with millimeter (mm) and sub-mm wavelengths (1-10 MHz ultrasound frequency). As a comparison, 2 MHz ultrasound has a wavelength of 0.75 mm while 2 MHz electromagnetic waves have a wavelength of 2.5 cm; this improves energy coupling efficiency to miniaturized implants^22^. In addition, compared to EM, US experiences lower attenuation in soft tissues (e.g., US: ∼1-2 dB·cm^-1^ at 2 MHz versus EM: ∼10-12 dB·cm^-1^ at 2 GHz)^23^ and consequently has a higher US Food and Drug Administration (FDA) regulatory limit for power flux density (US: 720 mW/cm^2^ versus EM: 10 mW/cm^2^)^24^. Note that unlike EM, US power transfer through bone is more challenging since the US attenuation in bone is more than an order of magnitude higher than in soft tissue, and reflections and scattering occur at the soft tissue - bone interface^25^. Phased arrays have also been used to focus US energy to an implant in the body^26^, but for efficient US energy coupling, the external US transmitter needs direct contact with the tissue through ultrasound gel or a gel pad.

During the last decade, implantable systems based on ultrasonic mm-sized implants have demonstrated *in vivo* electrical neural recording^20^ from and stimulation^21^ of the sciatic nerve in an anesthetized rat model, *in vivo* tumor oxygenation in an anesthetized mouse pancreatic tumor model^27^, photodynamic therapy of tumor in an anesthetized mouse model^28^ and *in vitro* monitoring of physiological parameters such as temperature^29^ and pressure^30^. Here, we present a system for direct tissue O_2_ monitoring that integrates ultrasound (US) technology with a luminescence sensor, demonstrating, for the first time to our knowledge, continuous real-time in vivo O_2_ measurements at centimeter-scale depths in a clinically-relevant, anesthetized sheep (large animal) model and operation at great depths (≥ 5 cm) through *ex vivo* porcine (anatomically heterogeneous) tissue. The system described here avoids all of the drawbacks present in current O_2_-sensing technologies, including O_2_ consumption, susceptibility to biofouling, wired connections, large volume for implants, long readout time, and inability to operate in deep tissue. The system employs a fully implantable, wireless, battery-free O_2_ sensor that incorporates a single piezoelectric ceramic and a luminescence sensor, consisting of a µLED, an O_2_-sensing film, an optical filter, and a custom state-of-the-art integrated circuit (IC) fabricated in a 65 nm low-power CMOS process. The luminescence sensor operates on the same readout principle, phase luminometry, as most luminescent sensors^31, 32^ and probes^33, 34^, but with much lower power consumption and competitive or better O_2_ resolution than these sensors and probes. This is achieved by compact integration of the sensor components and a custom IC. Wireless power and bi-directional data transfer are enabled using a single US link from an external transceiver. The use of a custom single-link protocol, combined with the tight integration of wireless sensor components, results in a very small sensor volume (4.5 mm^3^). The small volume, the lightweight construction (∼17.4 mg), and the use of encapsulation materials enable implantation and provide potential capabilities for use with minimal tissue damage. The operation of the system is demonstrated *in vitro* in distilled water, phosphate-buffered saline (PBS) and undiluted human serum, *ex vivo* through porcine tissue, and *in vivo* in an anesthetized sheep model. The ability to monitor tissue oxygenation during physiological states *in vivo* is confirmed via surgical implantation deep under the biceps femoris muscle.

## Results

### A custom integrated circuit and a single ultrasonic link enabled a complete wireless optical sensor in a 4.5 mm^3^ volume

An external ultrasonic transceiver directed ultrasound (US) energy from outside of the body toward an implant placed in muscle or deep tissue (Fig. 1a). This wireless acoustic link provided power to the sensor and allowed bidirectional data transmission. A custom mixed-signal benchtop system drove the external transducer (see Fig. 3a, left-top and Methods). The implant consisted of a single piezoelectric crystal for US energy harvesting and data communication coupled to a custom luminescent optical sensor for dissolved O_2_ detection.

**Fig. 1.**
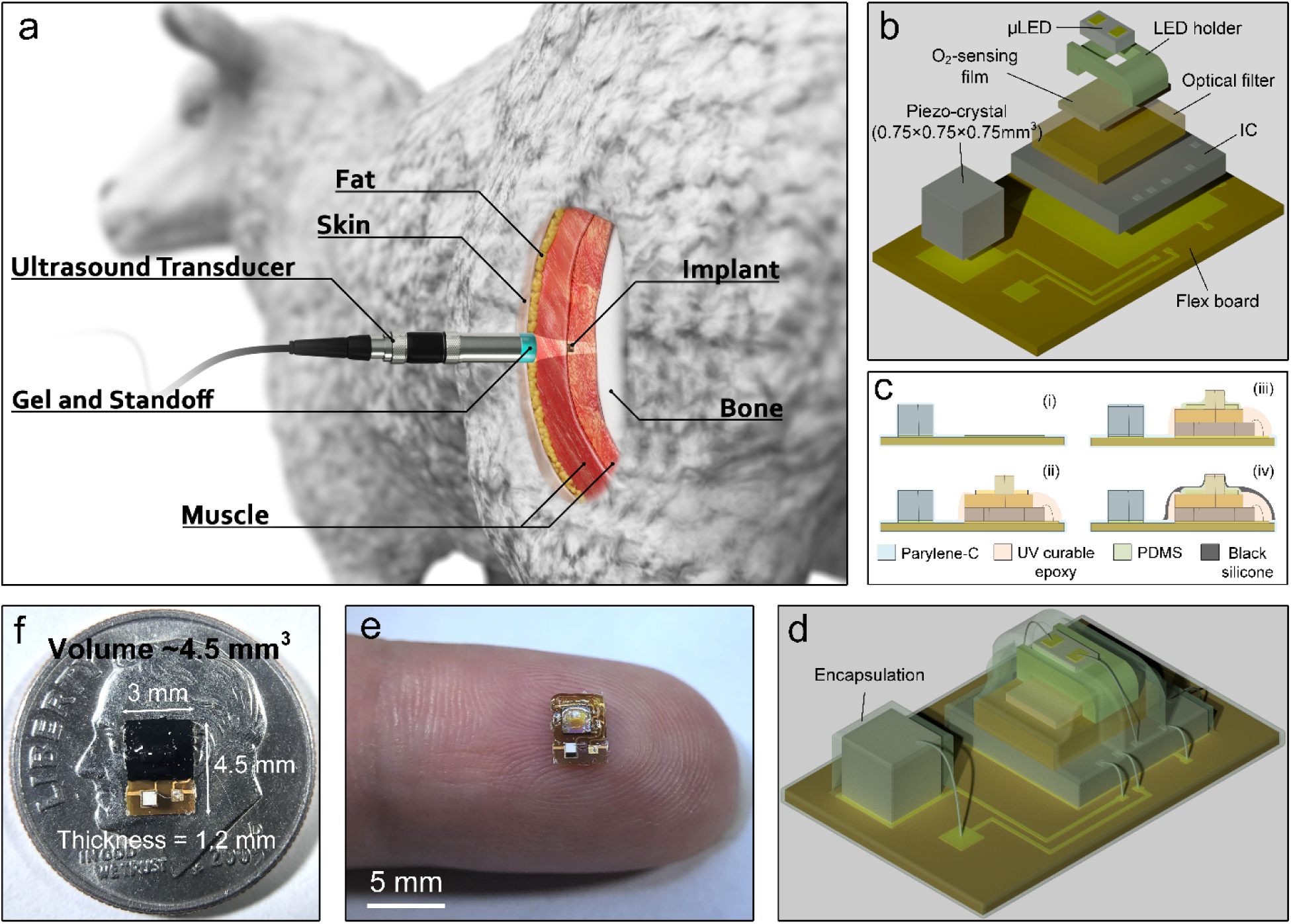
Wireless O_2_ monitoring system overview. **a**, Schematic demonstration of the system, as demonstrated in this paper for local tissue O_2_ monitoring in sheep. The free-floating, wireless O_2_ implant was surgically placed underneath the biceps femoris muscle, and the wound was closed with surgical sutures. The external ultrasound transducer, placed on top of the closed surgical site, established a wireless acoustic link to the implant. The external transducer powered the implanted sensor by delivering acoustic energy through the tissue and then listened for backscatter reflections from the sensor’s piezoelectric crystal, in which digital sensor data was encoded. The external transducer was driven by a custom mixed-signal system, including decoding and storage of the wireless O_2_ data received by the external transducer. **b**, An expanded view of the wireless sensor platform including a piezo crystal and a luminescence sensor, consisting of a µLED, a 3D- printed µLED holder, an O_2_-sensing film, an optical filter, and an IC. **c**, Wireless sensor fabrication steps: (i) A piezo-crystal was bonded with conductive silver epoxy, wire-bonded to a flexible PCB and encapsulated with parylene-C; (ii) other sensor components were assembled on the flex board, and only the regions where the wire-bonds are located were encapsulated with UV curable epoxy; (iii) small gap (∼50 µm) between the film and the µLED holder was filled by PDMS; (iv) sensor parts excluding the piezo-crystal were encapsulated with black silicone. **d, e**, Schematic diagram (**d**) and photograph (**e**) of the wireless sensor before black silicone encapsulation and size comparison to a finger (**e**). **f**, Top view of the 4.5 mm^3^ fully-packaged wireless sensor: piezo crystal at bottom, luminescent sensor at top, and size comparison to US dime.

### The entire implant was manufactured with standard medical implant processes

The implantable system (Fig. 1b) consisted of a 750×750×750 µm^3^ lead zirconate titanate (PZT) piezoelectric crystal for wireless power harvesting and data communication, a µLED with an emission peak at ∼465 nm for optical excitation, a thin film for encapsulation of O_2_-sensing luminescent dyes, an optical filter for excitation light rejection, a 3D-printed µLED holder for mechanical support during wire bonding of the µLED, and an IC with a size of 1.96×1.96 mm^2^ (Supplementary Fig. 1). To assemble the sensor (Fig. 1c), a piezo crystal was attached with conductive silver epoxy and then encapsulated with ∼10 µm-thick Parylene-C film, insulating the crystal from the environment; remaining components were then assembled on the flex board. Wire-bonds were encapsulated with ultraviolet (UV) curable epoxy, chosen due to its high bond strength^35^. Finally, all components (excluding the piezo) were coated with a ∼180 µm-thick layer of highly O_2_-permeable black silicone, acting as an optical isolation, to avoid background interferences by the luminescence of tissue or blood^36^. The fully-packaged sensor (Fig. 1d-f) measured 3 mm×4.5 mm×1.2 mm, occupied 4.5 ± 0.5 mm^3^ volume, and had a detection volume of ∼0.26 mm^3^ (estimated from the material volume where O_2_ molecules diffuse through to the O_2_-sensing film under the µLED). The resonant frequency of the piezo determined the carrier frequency of the ultrasound link; as this frequency was set by the crystal thickness and aspect ratio ^37^, crystal geometry was chosen to maintain a reasonable tradeoff between the frequency-dependent acoustic loss in tissue^38^, the capacity of power harvesting, and impact on total implant size.

### A complete biocompatible phase luminometric oxygen optrode was built into the implant

The biocompatible oxygen sensing film was polydimethylsiloxane (PDMS) containing 10 µm- diameter silica particles with surface-adsorbed ruthenium (Ru) dyes^39, 40^ (Fig. 2a, inset). The film thickness (∼100 µm) and the amount of silica particles in PDMS (∼8.3%) were adjusted to maintain a reasonable tradeoff between luminescence intensity, emitted from Ru-dyes under blue-light excitation, and O_2_ response time^39, 41, 42^. PDMS was used as polymer matrix due to its excellent O_2_-permeability, biocompatibility, optical transparency, and high solubility for the fillers, preventing agglomeration^43^. The Ru(dpp)_3_(ClO_4_)_2_ complex was selected as an O_2_-sensitive dye, mainly due to its large Stokes shift, relatively long excited-state lifetimes, and high photostability^36^.

**Fig. 2.**
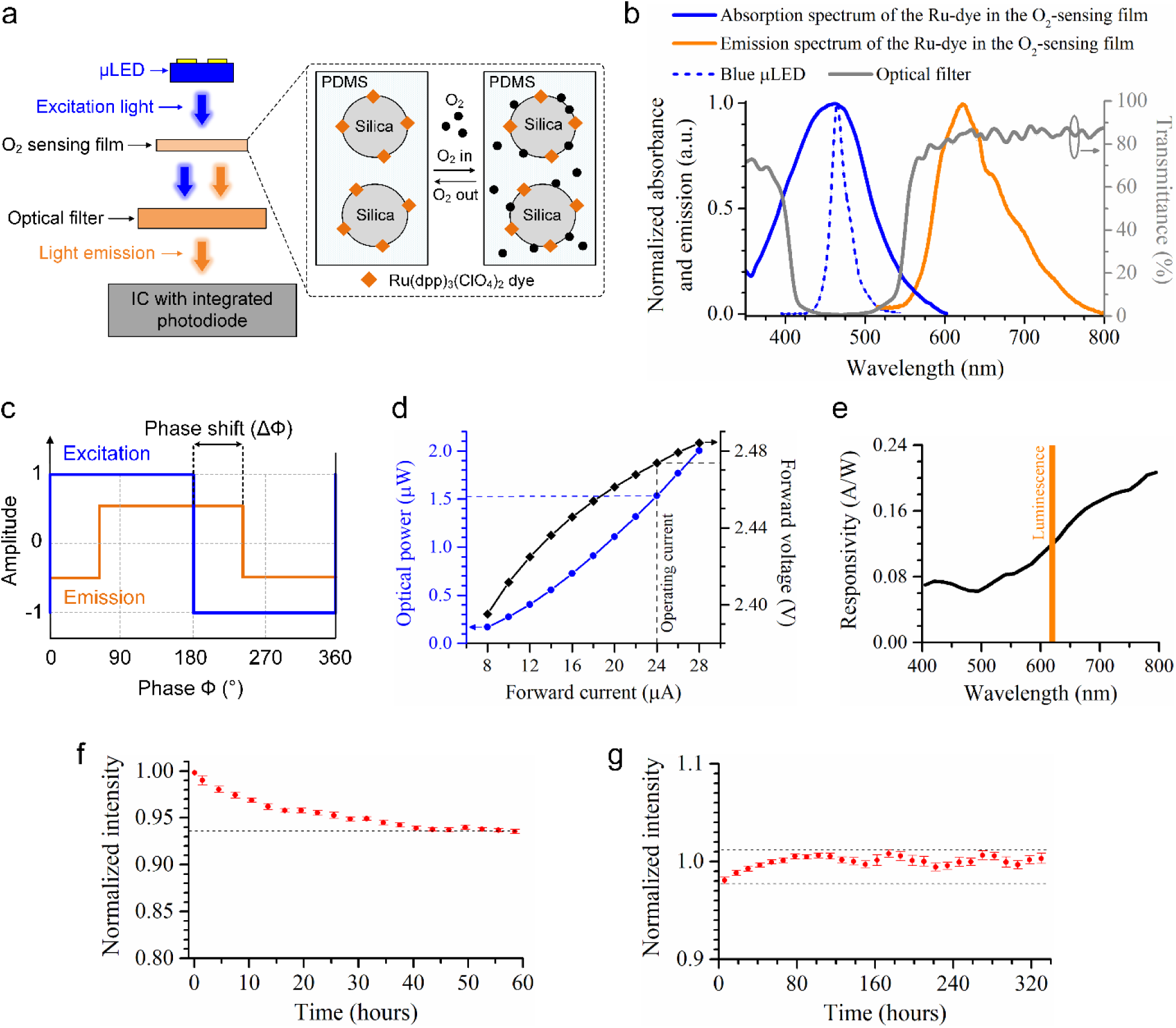
Biocompatible O_2_-sensing film, operating principle of the luminescence O_2_ sensor, and its optical characterization. **a**, An expanded cross-sectional view of the luminescence O_2_ sensor. **inset**, Model for the locus of Ru-dyes and O_2_ molecules in silica-containing PDMS. The orange squares and black circles represent the Ru-dyes and O_2_ molecules, respectively. The Ru-adsorbed silica particles were dispersed in PDMS. **b**, Normalized absorption and emission spectra of the Ru-dye in the O_2_-sensing film along with normalized emission spectrum of the blue µLED and transmission spectrum of the optical filter. **a, b,** In operation, the blue µLED, placed on the sensor platform to illuminate the O_2_-sensing film, produces light with a peak intensity at ∼465 nm that excites the Ru-dyes in the O_2_-sensing film. The excited Ru-dyes emit luminescence with a peak at ∼621 nm. A long-pass optical filter with a ∼550 nm cut-on wavelength suppresses excitation light and transmits luminescence, enabling the Ru-dye emission to be detected by an integrated circuit (IC) with an integrated photodiode. **c**, Illustration of frequency-domain luminescence excitation and emission signals. **d,** Current-voltage-light output characteristics of the blue µLED. **e,** Responsivity spectrum of an integrated photodiode with a 300×300 µm^2^ active area and a reverse bias voltage of 0.6 V. The orange shaded area highlights wavelength ranges with strong luminescence emission from the excited Ru-dyes. **f,** Photobleaching of Ru-dye in the O_2_-sensing film under continuous square-wave illumination with a peak excitation light power of ∼1.53 µW at the operating forward current of 24 µA, resulting in an average optical power density of ∼4.9 µW/mm^2^ at the surface of the film, in air (21% O_2_) at room temperature for a period of 60 h. **g,** Normalized luminescence intensity of Ru-dye in the PDMS film as a function of time after immersion of the same fully-packaged O_2_ sensor used for the photobleaching test in PBS solution at 37 °C in room air. During the test, the sensor was operated with the same operating conditions as in the photobleaching test.

**Fig. 3.**
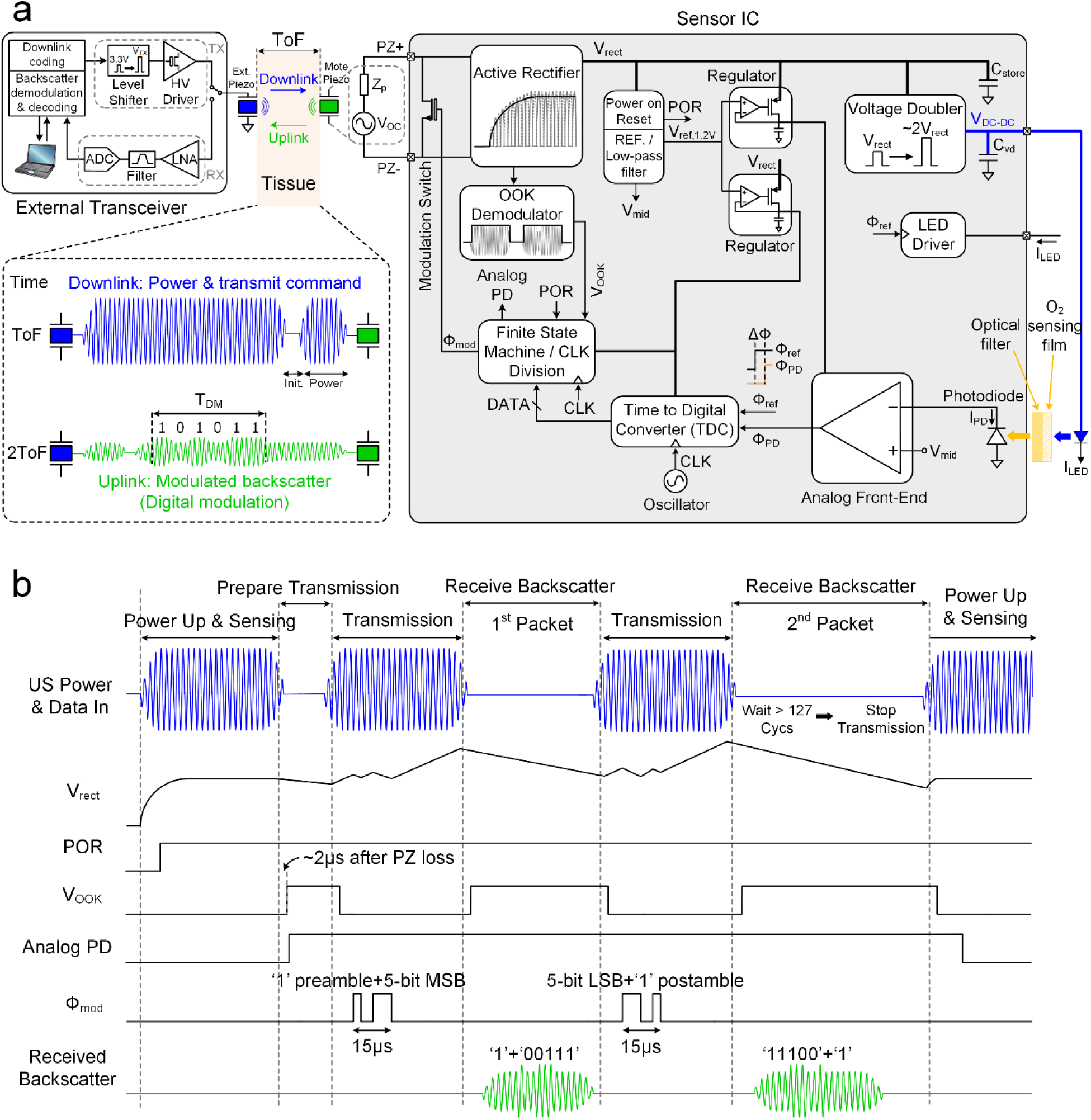
Entirely wireless oxygen monitoring system block diagram. **a**, The external transceiver is shown on the left-top, the ultrasound (US) link in the middle-top, and the wireless sensor on the right. **left-top**, The external transceiver consists of transmit (TX) and receive (RX) paths, where the TX path encoded downlink data onto a 2 MHz carrier. During TX operation, a level-shifter boosted a low-voltage transmit signal from a digital controller, and a high-voltage pulser drove an external piezo transducer. The RX path was enabled when the TX path was disabled. Reflected US backscatter from the sensor’s piezo crystal was captured by the same external piezo transducer, which was digitized by the RX chain. **middle-top**, The external piezo coupled to the outside surface of tissue produced US waves traveling through tissue; these arrived at the sensor after one time-of-flight (ToF). The downlink provided power and a transmit command for the sensor. The uplink consisted of amplitude-modulated backscattered US waves that arrived at the external piezo 2ToF after being sent during TX. **right**, The sensor IC architecture and **b**, timing diagram. The IC rectified electrical power harvested by the sensor’s piezo, and the power-on-reset (POR) initialized the O_2_-sensing operation. The OOK demodulator detected the downlink US envelope, producing a notch. The first notch started uplink transmission. Two data packets were transmitted via digital amplitude modulation of the US backscatter; the first packet contains five MSBs and one-bit preamble. The uplink data was received by the external piezo when the external transceiver was switched to RX. The uplink transmission was stopped when the third notch duration was longer than ∼64 µs (127 oscillations of a 2 MHz carrier). The IC circuitry was duty-cycled during uplink transmission to reduce energy consumption.

Upon light excitation at ∼465 nm, the Ru-dyes emit light with a peak intensity at ∼621 nm, enabling excitation light to be suppressed through an optical filter (Fig. 2a,b). The excited Ru- dyes undergo collisional quenching with O_2_-molecules, leading to a reduction in luminescence intensity and lifetime. Note that collisional quenching is a photo-physical process, enabling reversible O_2_ sensing capability^36^. Both the intensity (Ι) and lifetime (τ) depend on O_2_ concentration according to the Stern-Volmer equation^32^: I_0_/I = τ_0_/τ = 1+K_SV_·[O_2_], where I_0_ and τ_0_ (∼6.4 µs) are the intensity and lifetime at zero O_2_ and K_SV_ is the Stern-Volmer constant. Either intensity or lifetime can be measured to compute dissolved O_2_; however, luminescent lifetime (τ) is independent of variations in light source intensity and dye concentration, inner filter effects, and photobleaching (to a wide extent), all of which are main limitations of intensity-based sensors^34, 44, 45^.

This sensor was designed to operate on the principle of *phase luminometry*, wherein the phase shift (ΔФ) between the excitation and emission light was measured to determine the luminescent lifetime (τ). During operation, the emitted light was square-wave modulated at a fixed operating frequency (*f_op_* = 20 kHz), exciting the Ru-dyes in the PDMS film with a peak excitation power of ∼1.53 µW, resulting in an average power intensity of ∼4.9 µW/mm^2^ at the film surface (Fig. 2c, d). The excited Ru-dyes produce emission with a typical average power density of ∼8 nW/mm^2^ at 37 °C, ∼160 mmHg (room air) O_2_ concentration and the same *f_op_*, but with a phase shift (ΔФ) relative to the phase of the excitation light (Fig. 2c). The emission was detected by a 0.6 V reverse-biased, on-chip nwell/psub photodiode with an active area of 300×300 µm^2^ and a responsivity of ∼0.12 A/W at the peak emission wavelength of ∼621 nm after filtering the excitation light using a long-pass optical filter (Fig. 2a,e). The resulting phase shift (ΔФ) equals tan^-1^(2π*f_op_*τ)≈ω_op_τ for ω_op_τ << 1 and is directly dependent on the luminescence τ that, in turn, is related to local O_2_ concentration via the Stern-Volmer equation.

Photobleaching of the Ru-dye was evaluated using a fully-packaged O_2_ sensor (Fig. 1f) continuously operated in room air (21% O_2_) at room temperature for a total period of 60 h. The luminescence intensity decreased rapidly to ∼96.8% of its initial value in the first 10 h and at a slower rate from ∼96.8% to ∼93.6% in the next 50 h, indicating that the sensor could be operated with a duty cycle of 1%, corresponding to 14.4 min continuous operation in a single day, for 250 days with an only ∼6.4% drop in the luminescence intensity. After the bleaching test, a long-term continuous test (14 days) of the same sensor immersed in phosphate-buffered saline (PBS, 1×) solution at 37 °C was conducted in room air to evaluate dye leaching from the sensing film; the fluctuation in the luminesce intensity was within ±1.7% and did not exhibit a decreasing trend in 14 days.

### A custom integrated circuit drove sensor operation and communication for < 150 µW average power consumption

A functional block diagram of the system is shown in Fig. 3 (see reference^46^ for details of the sensor IC). During operation, the controller driving the external transducer alternated between transmit (TX) and receive (RX) modes. When in TX mode, it drove the transducer with a 2 MHz carrier wave onto which it encoded digital information in discrete pulses (Fig. 3a, inset, blue). The implant’s piezo crystal began harvesting energy upon the arrival of an ultrasound (US) pulse; this energy was rectified and stored on the on-chip storage capacitor (C_store_ = 2.5 nF) by an active full-wave rectifier. The energy in the cap allowed the implant to operate during the period when the external transducer was in RX mode. The rectified voltage (V_rect_) was regulated by the low-dropout (LDO) regulators, providing a constant 1.2 V for powering other circuits. When the LDO voltages exceeded a predetermined level (∼1 V) during the power-up period, a power-on-reset (POR) signal was triggered to initialize the IC. During an initialization period (∼150 µs), the rectified voltage was boosted by a voltage doubler to turn on the µLED. After the initialization period, an LED driver drove the µLED with a 24 µA, 20 kHz square-wave current, starting the O_2_-sensing operation. The µLED excited the O_2_-sensitive luminescent dyes, producing luminescence. The luminescence light, after suppressing the excitation light through an optical filter, was detected by an on-chip photodiode. The light-induced photocurrent (I_PD_) was converted to a voltage, and the resulting AC voltage was compared to its DC component by an analog front-end (AFE) for zero-crossing detection^47^ (Supplementary Fig. 2), generating a time-delayed luminescence signal. A time-to-digital converter (TDC), operating with a 16 MHz on-chip clock generated by a 5-stage current-starved ring oscillator_48_, converted the time delay (phase difference, ΔΦ) between the reference signal (Φ_ref_), used to drive the μLED, and the luminescence signal (Φ_PD_) into a 10-bit digital data. The 10-bit data was serialized and divided into two equal 15 µs-long data packets with a preamble and a postamble by a finite state machine; the first packet contains the most significant bits (MSBs). The measured minimum detectable average optical power, yielding a signal-to-noise ratio (SNR) equal to 1 at a 1 Hz bandwidth^49^, was ∼1.3 pW at the peak emission wavelength of ∼621 nm and the operating frequency of 20 kHz for the optical readout, which was dominated by the noise of the AFE. The SNR was ∼53 dB under a typical ∼6.7 nW/mm^2^ light power after the optical filter, produced by the excited Ru-dyes in the O_2_-sensing film, for the sensor operated in room air at 37 °C.

An uplink data transmission began when the on-off-keying (OOK) demodulator detected a “falling edge” in the US data input from the external transceiver, generating a notch (V_OOK_). The notch served as a reference to time synchronize the sensor IC and the external transceiver during uplink transmission; a data packet was transmitted to the external transceiver after a notch with a duration of shorter than ∼64 µs, equivalent to 127 oscillations of a 2 MHz US carrier. Data packets were encoded in the US reflections from the sensor’s piezo and transmitted via digital amplitude modulation of the backscatter. Backscatter amplitude modulation was achieved by modulating the electrical load impedance (R_load_), in shunt with the piezo impedance (Z_p_), through a modulation (transistor) switch controlled by Ф_mod_, changing the US reflection coefficient at the piezo boundary and thus the amplitude of the backscatter^50, 51^. When the transistor switch was turned on to transmit the O_2_ data, the R_load_ across the piezo was reduced from a resistance value higher than 80 kΩ, depending on the IC power consumption and the amplitude of the rectifier input voltages, to ∼0.5 kΩ (transistor switch on-resistance). The uplink transmission stopped when the notch duration was kept longer than ∼64 µs.

The total area of the IC die, fabricated in a TSMC 65 nm low-power CMOS process, is ∼3.84 mm^2^ (Supplementary Fig. 1). The minimum electrical input power required for proper operation of the IC was ∼150 µW, generating a rectifier voltage (V_rect_) of ∼1.36 V, during the O_2_-sensing phase. Power-intensive circuits (AFE, LED driver, voltage doubler, and TDC) were duty-cycled off during uplink transmission, reducing IC power consumption to ∼22 µW and thus avoiding the need for a large off-chip C_store_. The average power dissipation of the IC drops to less than 150 µW, including the rectifier’s power conversion efficiency, during operation, depending on the O_2_ sampling rate. The sampling rate (*f_s_*) of the system was externally controlled through the external receiver.

During operation, the external transceiver was switched from TX to RX mode to capture uplink data encoded in the backscatter reflections from the sensor’s piezo. The RX path demodulated and decoded the received backscatter, generating real-time O_2_ data. The data was sent to a computer through a serial link for data storage and further analysis. In order to avoid overlapping TX and RX pulses, the data packet duration (T_DM_) was kept shorter than the round-trip time-of-flight (2ToF) of a US pulse between the sensor’s piezo and the external transducer (Fig. 3a, inset, green), limiting the minimum operating distance between the sensor and the external transducer. Note that, in order to overcome this limitation, the digital data was divided into two parts that were subsequently sent to the external transceiver. Supplementary Fig. 3 shows an alternative communication protocol for sensors implanted deeper than 5 cm because of the longer ToF to these depths. Compared to the protocol in Fig. 3b, the alternative protocol reduces the time spent during data transmission and hence increases the sampling.

### The system sampled oxygen 350 times per second with a resolution of < 5.8 mmHg/**√**Hz across the physiologically relevant oxygen range of interest (0-100 mmHg) and a bit error rate of < 10^-5^

The system was first characterized in a water tank setup (Fig. 4a and Supplementary Fig. 16) where the distilled (DI) water temperature was kept constant at 37 ± 0.1 °C to simulate physiological temperature, and O_2_ concentration was monitored via a commercial O_2_ probe and varied by controlling the ratio of O_2_ and nitrogen (N_2_) supplied to the water tank. Distilled water has an acoustic impedance similar to soft tissue (∼1.5 MRayls)^52^.

**Fig. 4.**
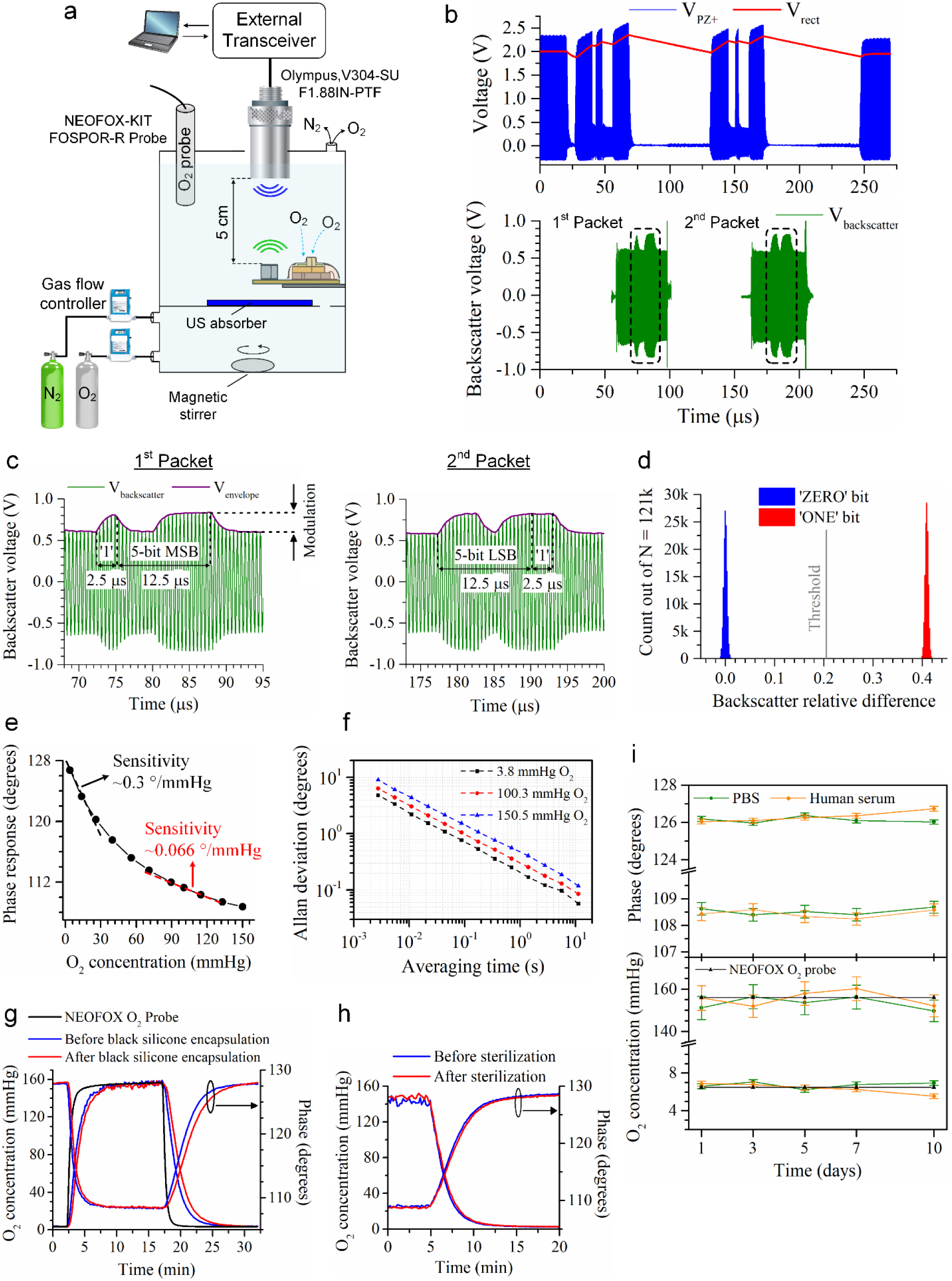
*In vitro* characterization of the wireless oxygen-sensing system. **a**, Wireless sensor operation in distilled (DI) water. The depth was set to 5 cm, and water O_2_ levels were precisely controlled using two identical gas flow controllers and a reference O_2_ sensing probe (NEOFOX) during measurements. **b**, Sensor waveforms and backscatter signal were recorded in the wireless measurement of a single O_2_ sample. **c,** Zoomed-in data packets show digital amplitude-modulation of the US backscatter. **d**, Backscatter relative difference for 121k O_2_ samples, showing ∼41% modulation depth. To detect each bit that is either “0” or “1”, the amplitude of the backscatter signal in the data packets was compared to a threshold. The wireless system achieved an uplink bit error rate (BER) less than 10^-5^ (0 out of 121k samples) in the measurement, demonstrating a robust data uplink. **e**, System response to various O_2_ concentrations. The measurement was performed at 350 samples per second *f_s_* and 5 cm depth in the O_2_ range of 3.8-150 mmHg. **f**, Allan deviation of the raw data (shown in Supplementary Fig. 5a). **g, h,** O_2_ sensor response to changes in O_2_ concentration in DI water at 37 °C before and after (**g**) black silicone encapsulation and (**h**) ethylene oxide (EtO) sterilization. **i,** Data from the O_2_ sensors incubated in PBS and undiluted human serum at 37 °C for 10 days.

The system was operated as described above, with an implant placed at a depth of 5 cm. Measured waveforms and backscatter signal were recorded for the system operated at a sampling rate (*f_s_*) of 350 samples per second (Hz) using an acoustic field with a spatial-peak time-average intensity (I_SPTA_) of 237 mW/cm^2^ (∼32.9% of the FDA safety limit, I_SPTA_ derated = 720 mW/cm^2^, for diagnostic ultrasound) (Fig. 4b). The system exhibited a power transfer efficiency from the acoustic power at the surface of the implant’s piezo crystal to the electrical input power of the IC of ∼20.4% and a US link power transfer efficiency of ∼3.9%, defined as the ratio of the electrical input power of the IC to the acoustic power emitted from the external transducer. Each data packet was 15 µs long, containing 6-bits with 2.5 µs duration (Fig. 4c). The first data packet began with a ‘1’ preamble followed by 5-bit MSBs, and the second data packet began with 5-bit least significant bits (LSBs) followed by a ‘1’ postamble. The system achieved a modulation depth of ∼41%, and an uplink bit error rate (BER) less than 10^−5^ (0 out of 121k samples) with the optimal threshold, minimizing BER, determined by the external transceiver (Fig. 4d).

The system response to various O_2_ concentrations and the Allan deviation of the data are illustrated in Fig. 4e,f. An Allan deviation analysis^53^ was used to quantify the noise performance of the system. The system operated at 350 samples per second *f_s_* exhibited an O_2_ sensitivity greater than ∼0.066 °/mmHg and a phase (Φ) resolution less than 0.38°/√Hz, yielding an O_2_-resolution better than 5.8 mmHg/√Hz across the relevant O_2_ range of interest (< 100 mmHg). Since the Ф resolution in the system is dominated by jitter noise of the AFE and the luminescence intensity (I) increases with a decreasing O_2_ concentration, the system SNR and hence Ф-resolution improves for lower O_2_ levels. The nonlinearity of the phase readout circuit, including the photodiode, AFE, and TDC, was characterized using a function generator that produced a variable Φ-shifted signal to drive the μLED. The worst-case nonlinearity, computed using the endpoint method, was less than 0.27 LSB = ∼0.12° (see Supplementary Fig. 4a). The τ- based Stern-Volmer plot (Supplementary Fig. 5b), obtained using the equation: ΔФ = tan ^1^(2π*f_op_*τ), reveals the nonlinearity at the system output, which is mainly due to heterogeneous dispersion of silica particles in the O_2_-sensing film^54^.

The luminescence O_2_ sensor response to changes in O_2_ concentration is reversible (Fig. 4g). Note that we used a calibration curve and equation to convert sensor phase output to O_2_ concentration (partial pressure of oxygen, pO_2_) in mmHg in this study (see Supplementary Fig. 6). The response time (the time required to reach 90% of the steady-state value) of the sensor before black silicone encapsulation from ∼3.5 to ∼156 mmHg O_2_ and vice versa were ∼210 s and ∼257 s, respectively; this increased to ∼250 s and ∼320 s after black silicone encapsulation of the O_2_ sensor.

In order to assess the potential effect of sterilization on the O_2_ sensor functionality, the sensors were first sterilized in ethylene oxide (EtO), and then their response to O_2_ changes was tested in DI water at 37 °C. The sensor response to O_2_ variation before and after sterilization was nearly identical (Fig. 4h), indicating that these sensors are sterilizable without loss of their functionality.

To evaluate the ability of the O_2_ sensor to resist *in vivo* biofouling, we conducted *in vitro* experiments in phosphate-buffered saline (PBS, 1×) and undiluted pooled human serum. Note that human serum, a complex fluid containing hundreds of different proteins^55^, was chosen to simulate *in vivo* fouling since nonspecific protein adsorption (fouling) on the implant surface is considered the initial step in triggering a foreign-body reaction^56^ and one of the critical factors causing failure of many implants^57^. The response of the O_2_ sensors incubated in PBS and serum at 37 °C was measured at low (6.5 mmHg) and high (156 mmHg) oxygen levels and different times during 10-days (Fig. 4i). The sensors demonstrated no apparent loss of sensitivity to O_2_ changes during these 10-days.

### The wireless system accurately reflected tissue oxygenation (pO_2_) in a physiologically relevant large animal model

We next tested the clinical utility of the wireless, direct O_2_ monitoring system in a physiologically relevant large animal model. The sheep model is a standard in fetal, neonatal and adult disease states due to the remarkable similarities in cardiovascular and pulmonary physiology^58, 59^, neurobiology^60^, as well as metabolism^61^. Anesthetized juvenile or adult sheep (n = 2 animals) were intubated and mechanically ventilated. The biceps femoris was carefully dissected and the wireless sensor, as well as a commercial wired pO_2_ sensor (NEOFOX), were placed in the plane below the muscle layer and the muscle, as well as overlying skin, were carefully closed above (Fig. 5a-c and Supplementary Fig. 20a,b). The ultrasound transducer, attached to a five-axis micromanipulator for fine alignment, was placed on top of the skin layer on an acoustic standoff pad (Fig. 5d and Supplementary Fig. 20c).

**Fig. 5.**
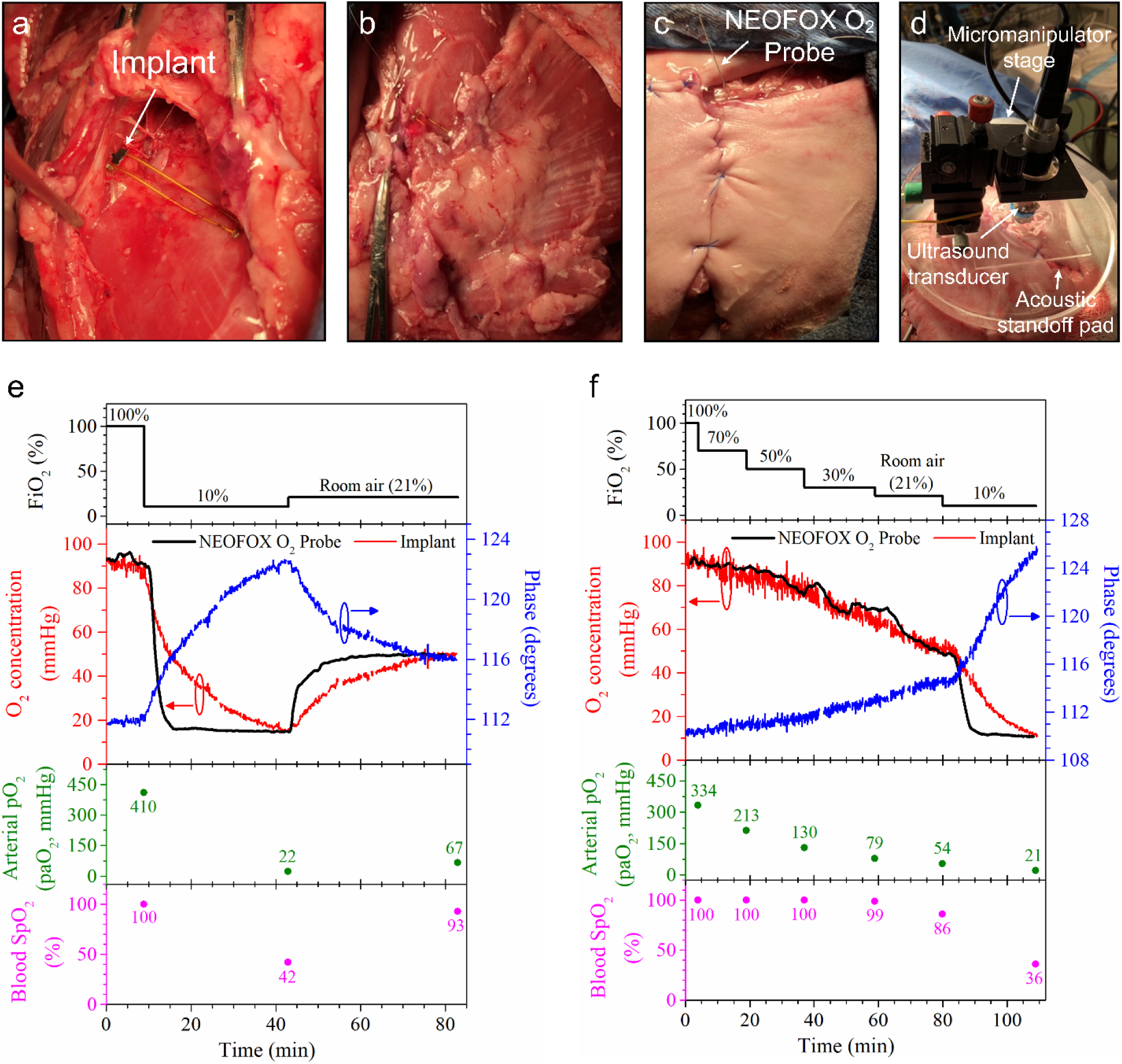
Surgical placement of and recording from the implantable, wireless O_2_ sensor under conditions of normoxia, hyperoxia, and hypoxia. **a,** The wireless sensor was placed underneath the biceps femoris muscle in anesthetized sheep following careful surgical dissection. **b, c.** The biceps femoris muscle was layered over the sensor, completely covering it (**b**), and the overlying skin was sutured using an interrupted approach (**c**). **d,** The micromanipulator stage, housing the ultrasound transducer, was placed on top of the skin layer on an acoustic standoff pad**. e,** The anesthetized animal was provided with 100% inspired oxygen via an endotracheal tube, followed by a hypoxic gas mixture of 10% O_2_ achieved via nitrogen blending and confirmation with an inline O_2_ detector, followed by ventilation with room air (21% O_2_). Tissue O_2_ concentration readings were continuously monitored via the wireless O_2_ sensor as well as the wired commercial NEOFOX probe. Corresponding paO_2_, SpO_2_ and FiO_2_ readings are provided. **f,** Stepwise reduction in FiO_2_ resulted in corresponding stepwise reductions in tissue pO_2_ readings that showed excellent concordance between the wireless O_2_ sensor and the commercial probe. Corresponding paO_2_, SpO_2_ and FiO_2_ readings are provided. Note that for FiO_2_ values above room air, SpO_2_ readings are unable to approximate paO_2_ or tissue pO_2_ levels.

**Fig. 6.**
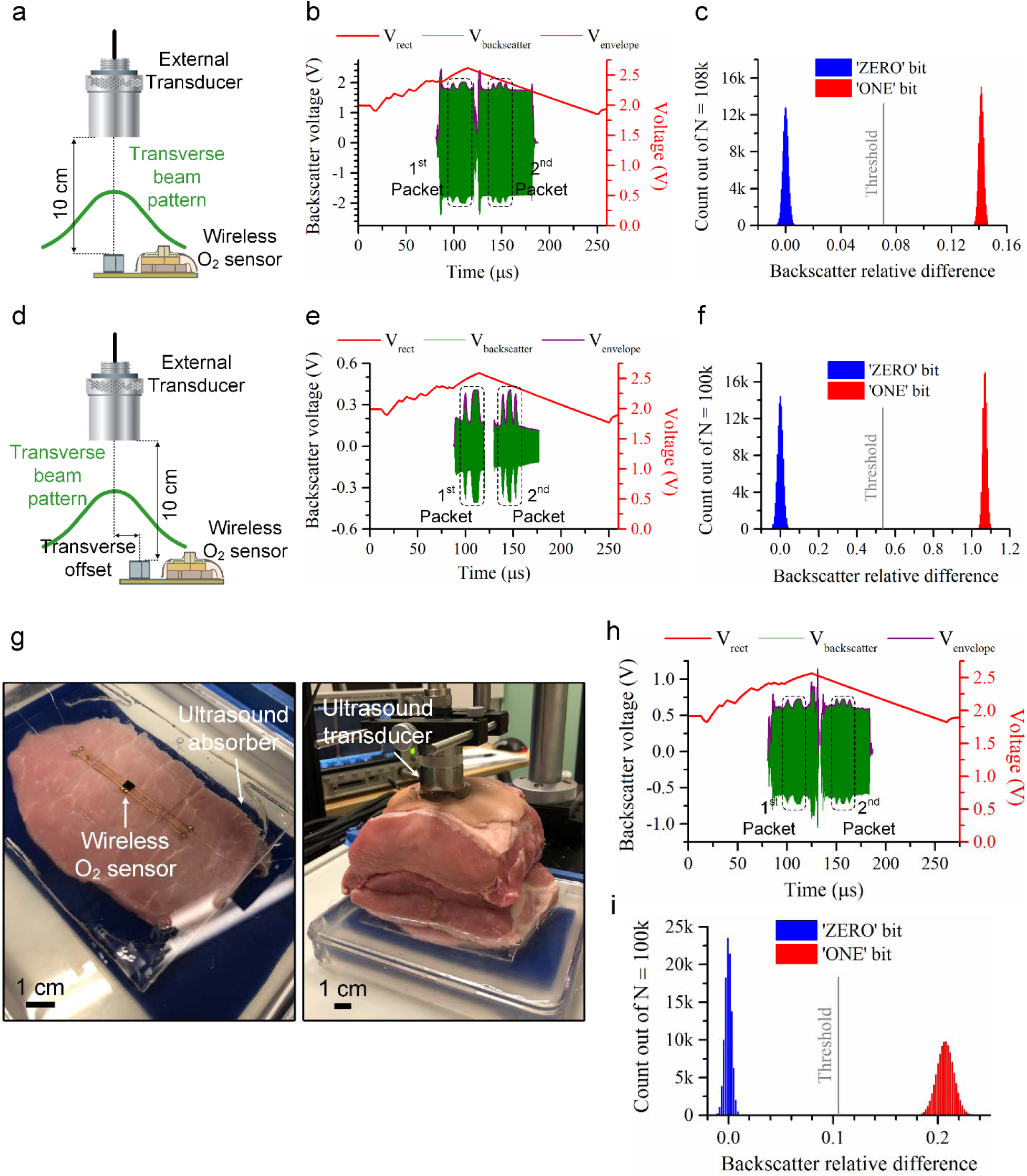
*In vitro* and *ex vivo* uplink characterization of the wireless oxygen-sensing system. **a, d,** Wireless sensor operation at 10 cm depth in distilled water without (**a**) and with (**d**) an intentional transverse offset of the central axis of the acoustic field to that of the sensor’s piezo. **b, e,** Sensor waveform and backscatter signal were captured in the wireless measurement of a single O_2_ sample for the sensor operated *in vitro* at 10 cm depth without (**b**) and with (**e**) a transverse misalignment. **c, f,** Backscatter relative difference for wirelessly recorded O_2_ samples ≥100k from the sensor operated in vitro at 10 cm depth without (**c**) and with (**f**) a transverse misalignment. **g,** The O_2_ sensor was wirelessly operated at a depth of 10 cm through an inhomogeneous sample of fresh, *ex vivo* porcine tissue, in which acoustic waves passed through approximately 3 mm ultrasound gel, 1 mm skin, 1 mm fat, and 95 mm muscle tissue. **h,** Sensor waveform and backscatter signal were recorded in the wireless measurement of a single O_2_ sample for the sensor operated at 10 cm depth through a porcine specimen. **i,** Backscatter relative difference for 100k O_2_ samples, showing ∼21% modulation depth, from the sensor operated at 10 cm depth through a porcine tissue specimen. The system achieved an uplink BER of < 10^-5^ in the measurement.

As seen from the data (from animal A) in Fig. 5e, and determined by the wired pO_2_ sensor, acutely adjusting the inspired O_2_ concentration from 100% to 10% resulted in a rapid reduction in muscle pO_2_ from approximately 90 mmHg to around 20 mmHg. The wireless sensor similarly was able to accurately reflect muscle pO_2_ levels.

More gradual, stepwise reductions in inspired oxygen content resulted in more gradual reductions in tissue pO_2_, which were accurately determined by our wireless sensor in real-time with similar kinetics as the commercial wired probe (Fig. 5f; the data from animal B). Importantly, simultaneous determination of blood hemoglobin saturation via pulse oximetry was not capable of detecting differences in tissue or blood oxygenation above room air (21%), as all hemoglobin is fully saturated beyond this point.

In sum, our mm-scale wireless implantable oxygen sensor accurately reflects tissue oxygenation status under physiological states and therefore has significant potential to augment clinical decision making in settings where tissue or patient oxygenation status warrants careful monitoring.

### The wireless system demonstrated robust wireless operation both *in vitro* and *ex vivo* with the implant operated at 10 cm depth

To further evaluate uplink performance, the system was operated at 350 Hz *f_s_* and 10 cm depth in DI water with and without intentional misalignment using the same water tank setup (Supplementary Fig. 16) (Fig. 6a,d) and through fresh, *ex vivo* porcine tissue (which presented inhomogeneous acoustic properties) (Fig. 6g). The system, while operating without any misalignment and using a 2 MHz acoustic field with an I_SPTA_ of 282 mW/cm^2^, exhibited a modulation depth of over 14%, a BER lower than 10^−5^ and a US link power transfer efficiency of ∼0.74% (Fig. 6a-c). When operated with an intentional transverse offset of ∼1.21 mm between the central axes of the acoustic field and the sensor piezo, the system demonstrated a modulation depth of ∼108% and a BER lower than 10^−5^, but at the expense of a ∼58.9 % increase in I_SPTA_, resulting in a concomitant reduction in the US link power transfer efficiency (Fig. 6d-f). As seen from the amplitude of the unmodulated backscatter signals (Fig. 6b,e), the increase in modulation depth was mainly due to reduced reflections from non-responsive regions (here, the sensor surface), indicating that an acoustic field with a smaller transverse beam spot size (i.e. comparable to the piezo size) can provide more robust uplink performance and higher US link power transfer efficiency.

In the *ex vivo* measurement, the external transducer generated US waves with 706 mW/cm^2^ I_SPTA_ de-rated by 0.3 dB·cm^-1^·MHz^-1^ (FDA-standard US attenuation in soft tissue^24^), producing an acoustic power of ∼228 mW at the transducer surface, that propagated through approximately 3 mm ultrasound gel, 1 mm skin, 1 mm fat, and 95 mm muscle tissue (Fig. 6g). The system achieved ∼21% modulation depth, a BER of < 10^-5^, demonstrating a robust uplink performance, and a US link power transfer efficiency of ∼0.08% with the sensor consuming ∼194 μW average electrical power (Fig. 6e,f). Note that uplink performance (modulation depth and BER) of the system depends on acoustic attenuation due to scattering and absorption in heterogeneous tissue, which varies for different tissue types/specimens. This is because acoustic absorption and dispersion may change the amplitude of the unmodulated backscatter signal received by the external transceiver during the time interval within the uplink data is received. Furthermore, US link power transfer efficiency is also a function of acoustic attenuation (see Methods); for example, the system operated through a porcine tissue specimen (Fig. 6g) exhibited a significantly lower link power transfer efficiency than the system operated in distilled water (Fig. 6a) since the US attenuation of a tissue sample is much higher than that of water.

## Discussion

This work presents the first miniaturization of a fully implantable optrode in the mm^3^ volume range, which is suitable for deep-tissue measurements. Although the focus of the study is a system for measuring oxygen tension *in vivo*, the fundamental technological achievement opens the door to minimally invasive pulse oximetric sensors^18^, pH sensors^62^, CO_2_ sensors^63^ among others^64–66^. Each of these, embodied in ultra-miniature and deep tissue systems, would open the door to novel diagnostics.

With regards to the measurement of deep-tissue oxygen tension, the application space of the wireless O_2_-sensing system is vast. Organ transplantation provides a clear example. The demand for organ transplantation continues to grow. In 2019, there were 94,863 candidates on the waiting list for renal transplants in the U.S. alone^67^. Monitoring graft oxygenation following organ transplantation is critical but typically relies on indirect methods that require skilled operators and provide only intermittent snapshots of tissue perfusion^68, 69^. Continuous and reliable monitoring of graft oxygenation following orthotopic liver transplantation, for example, may enable early detection of graft ischemia due either to hepatic artery thrombosis or graft vascular disease, allowing timely surgical re-exploration to minimize risk of graft loss, which could be fatal^2,70^. Importantly, these complications can occur months to years following transplant^3, 71^. Minimally invasive wireless modalities such as described here could enable real-time monitoring of graft oxygenation via wearable applications in the out of hospital setting, providing critical information regarding tissue oxygenation before the emergence of graft dysfunction, allowing timely intervention. This would additionally help differentiate parenchymal rejection from graft vascular disease when organ dysfunction emerges. Furthermore, more than 5.7 million patients are admitted annually to intensive care units (ICUs) in the United States (U.S.)^72, 73^. Assessment of tissue oxygenation is a fundamental need in this setting. Although the current iteration of our device requires surgical placement, future miniaturization efforts may enable semi-invasive/vascular approaches for probe placement. Depending upon the underlying pathology, the local oxygen supply-demand balance can be distorted during pathological states, such as observed during various forms of shock^74^. Thus, an inadequate delivery — for whatever reason — relative to demand will decrease tissue pO_2_. On the other hand, a primary reduction in metabolic demand or an inhibition or failure of mitochondrial oxidative phosphorylation will leave oxygen supply largely unaffected, and thus, the tissue pO_2_ may increase. A close matching of oxygen supply and demand, be it via an overall increase in delivery or decrease in turnover, will result in no net change in tissue pO_2_. Global measures of cardio-pulmonary performance such as cardiac output, oxygen delivery or blood pressure frequently do not reflect local metabolic demands at the organ and tissue level and can promote excessive fluid loading or inotrope dosing, worsening outcomes^75–77^. An important contributor in this setting is a lack of hemodynamic coherence between the microcirculation and the macrocirculation^78^. Given that these changes typically occur over minutes to hours, a slightly longer response time than typically observed for pulse oximetry would still yield important clinical information. Coupling direct measurements of the microcirculation^74^ with direct monitoring of tissue pO_2_ would greatly augment critical care management approaches. Precise measurements of tissue oxygen content are therefore instrumental for the proper management of shock states, but are currently limited to indirect or surface deep methods. Novel non-invasive as well as minimally invasive modalities for monitoring deep-tissue oxygenation as described here are clearly needed to advance our understanding and management of disease states where oxygen delivery or metabolism is compromised.

To clinically adopt the wireless O_2_-sensing system for the use of chronic, real-time *in vivo* O_2_ tracking, a number of technical challenges must be addressed. One of the main challenges is the post-surgical localization of the implant by an external transceiver since the post-surgical *in vivo* position may drift relative to any external fiducials; such movement can arise due to pressure from outside the body, movements or breathing of the subject, and scar formation. The *in vivo* localization before each pO_2_ measurement can be achieved with an external phased-array transceiver that utilizes ultrasound (US) backscatter information first to find and then track the time-dependent position of the implant in the body^79^.

A second challenge arises because acoustic attenuation due to scattering and absorption varies between different US propagation paths to the implant in heterogeneous tissue; a path with higher attenuation in tissue may significantly degrade power transfer efficiency and data transfer reliability of the system. For example, muscle tissue with a more unevenly distributed intramuscular fat content will exhibit greater acoustic attenuation^80^. Here too, an external transceiver with a large-aperture, multi-element transducer array capable of focusing US energy to the implant will allow for steering the US beam along a preferred path. Finally, a phased array could also potentially be used to interrogate multiple O_2_ sensors implanted in different locations of target tissue in a time-division multiplexing fashion or simultaneously^81^.

In addition to these improvements, chronic *in vivo* use of the wireless O_2_ sensor will require hermetic packaging to prevent biofluid penetration into the electronic sensor components (IC and μLED) and the piezo crystal. Traditionally such millimeter-scale implantable hermetic housings make use of ceramic or titanium enclosures brazed or microwelded to achieve the required hermeticity. This is an active area of work both commercially and academically; an extensive review was recently published^82^. Acoustic windows for efficient ultrasonic energy transfer into ceramic or metallic housings have recently been demonstrated in the academic literature^83^. For this work, we used biocompatible polymer materials (parylene-C, silicone and UV-curable epoxy to encapsulate the sensor given their ease of use for acute and semi-chronic experiments. Of course, it is well known that polymeric materials at these thicknesses are not suitable for long-term *in vivo* use of the implant due to their high water vapor permeability^84, 85^.

## Methods

### Fabrication of the oxygen-sensing film

The film fabrication included two steps. First, luminescent dyes, tris-(Bathophenanthroline) Ruthenium (II) Perchlorate (Ru(dpp)_3_(ClO_4_)_2_) (CAS 75213-31-9; GFS Chemicals), were immobilized on the surface of silica particles with a diameter of 10 µm (CAS 7631-86-9; LiChrosorb Si 100 (10µm); Sigma-Aldrich) at about 1:10 dye:particle ratio by weight. Briefly, 200 mg Ru(dpp)_3_(ClO_4_)_2_ complex was dissolved in 10ml ethanol (ACS reagent ≥99.5%, CAS 459844; Sigma-Aldrich). Silica gel was prepared by adding 2 g silica particles to 40 ml aqueous NaOH (0.01N; CAS 1310-73-2; Fisher Scientific) solution and magnetically stirring the mixture at a speed of 1000 rpm for 30 min. Next, the dye-containing ethanol solution was poured into the silica gel solution and stirred at 1000 rpm for 30 min. The dye-containing silica particles were filtered out of the solution through a filter with a pore size of 0.45 µm (Catalog number 165-0045; ThermoFisher Scientific), and then washed once in ethanol and three times in deionized water. All the supernatant was removed, and the dye-loaded silica particles were dried at 70 °C overnight.

Second, the dye-loaded silica particles were incorporated into polydimethylsiloxane (PDMS) to avoid problems related to dye leaching in aqueous media. 2 g dried silica particles were thoroughly mixed with 20 g PDMS prepolymer Part A and 2 g PDMS curing agent Part B (Sylgard 184; Dow Corning). A ∼100 µm-thick film was prepared by spinning a small amount of this mixture at 500 rpm on a microscope slide and then by curing it at 60 °C under dark and vacuum (< 10 Torr) for ∼7 days, to remove solvent and air bubbles. The cured film was kept under dark at room temperature for at least 24 h before use and stored under dark at room temperature.

### Design, fabrication, and assembly of the wireless oxygen sensor

The wireless sensor was built on a 100 µm-thick polyimide, flexible PCB with electroless nickel immersion gold (ENIG) coating (Rigiflex Technology). A 750 µm-thick lead zirconate titanate (PZT) sheet with a 12 µm-thick fired on silver electrodes was diced using a dicing saw with a 300 µm-thick ceramic- cutting blade. A 750 µm^3^ PZT cube was first attached to a flexible PCB using two-part conductive silver epoxy with 1:1 mix ratio (8331, MG Chemicals), and then the board was cured at 65 °C for 15 min, well below the PZT Curie temperature and the melting temperature of polyimide. The top electrode of the PZT was wire bonded to the PCB using a wedge bonder (747677E; West Bond) to create an electrical connection between the PZT and the IC. The board was then encapsulated with ∼10 μm-thick layer of parylene-C using chemical vapour deposition (Specialty Coating Systems) for insulation due to its biological inertness and resistance to a moisture^86^. The ∼10 μm-thick Parylene-C reduces the power harvesting efficiency of the PZT by ∼49% by damping its vibrations^21^. The metal pads on the PCB for the IC and its wire bonds were carefully exposed by scoring the parylene around the pads using a sharp probe-tip and removing the parylene layer. The IC was attached to the PCB using the same silver epoxy, cured at 65 °C for 15 min, and then wire bonded to the PCB. Next, a ∼250 µm-thick optical long-pass filter with a cut-on wavelength of 550 nm (Edmund optics) were attached to the top of the IC using medical-grade, UV-curable epoxy (OG142; Epotek). The same UV curable epoxy was also used to assemble other sensor components, including a µLED with dimensions of 650 µm×350 µm×200 µm (APG0603PBC; Kingbright) and its 3D-printed holder (Protolabs), to protect the wire bonds of the chip and µLED and provide insulation. After the assembly was completed, the ∼100 µm-thick O_2_-sensing film was slipped through the gap between the µLED holder and the optical filter. The small residual space between the µLED holder and the film was filled by PDMS (Sylgard 184; Dow Corning). PDMS Sylgard 184 Part A and B were mixed in the ratio of 10:1, degassed, poured between the space, and cured at room temperature for 48 h. Finally, the oxygen-sensing region on the IC was coated with a ∼180 µm-thick layer of biocompatible, highly O_2_-permeable black silicone. The black silicone consisted of two-part, low-viscosity silicone elastomer (MED4-4220, NuSil Technology, LLC) and black, single component masterbatch (Med-4900-2 NuSil Technology, LLC); the two silicone parts (A and B) were first mixed in a 1:1 weight ratio, and then the masterbatch (4% by weight) was added, thoroughly mixed, degassed for < ∼5 min, applied to the sensor surface, and cured at room temperature for 48 h.

In this work, PZT was selected as a piezoelectric material due to its high electromechanical coupling coefficient and high mechanical quality factor, providing high power harvesting efficiency. A lead-free biocompatible barium titanate (BaTiO_3_) ceramic with a slightly lower electromechanical coupling coefficient can be used in place of PZT^87^.

The volume of a wireless O_2_ sensor was measured by using a suspension technique^88^. In volume measurements, the sensor without test leads was suspended with a thin, rigid wire below the water surface in a container placed on an electronic balance with a measurement accuracy of 0.1 mg. The volume of the sensor was calculated from the weight difference of a water-filled container before and after submersion of the sensor in water; the weight difference, equal to the buoyant force, was divided by the density of water to determine the actual sensor volume. The volume measurements were performed using two separate sensors; the volume of each sensor was measured five times to determine reproducibility. The data obtained from all the volume measurements were presented by the mean and standard deviation values (mean ± 2s.d.).

### Optical characterization of components of the luminescence oxygen sensor

The absorption spectrum of the O_2_-sensing film and the transmission spectrum of the optical filter were measured with a Jenway 6300 spectrophotometer. The emission spectra of the sensing film and the blue µLED was measured using a fiber-coupled CCD spectrometer (Thorlabs, CCS200/M) operating at an integration time of 1 s and enabled with electric dark correction. The film samples were excited at 450 nm by a laser diode (Osram, PL450B, purchased from Thorlabs) driven with a Keithley 2400 source meter, and its emission was scanned in the range of 515-800 nm. The optical output power level of the µLED was measured using an optical power meter (Thorlabs, PM100D) equipped with a Si photodiode detector (Thorlabs, S121C). The current-voltage curve of the µLED was measured with a Keithley 2400 source meter. The responsivity of the photodiode as a function of wavelength was measured using a halogen lamp coupled to a monochromator, a reference photodiode (Thorlabs, FD11A Si photodiode) and an Agilent B2912A source meter. The same photodiode (FD11A) was also used to measure the output light intensity of the µLED.

### Photobleaching and leaching tests

Photobleaching of the Ru-dye in the O_2_-sensing film was evaluated using a fully-packaged O_2_ sensor (Fig. 1f) continuously operated in room air (21% O_2_) at room temperature for 60 h period. After the photobleaching test, the same sensor was continuously operated in phosphate-buffered saline (PBS, 1×) without calcium and magnesium (Corning; Mediatech Inc.) at 37 ± 0.1 °C in an oven (Test Equity Model 107) for 14 days, to assess leaching and further photobleaching of the Ru-dye in the film. In the tests, the sensor was electrically driven by differential, 2 MHz AC signals from a Keysight 33500B function generator, which are ac-coupled to the rectifier inputs of the IC. The output of the transimpedance amplifier (TIA) (Supplementary Fig. 2) was connected to a buffer (LTC6268; Linear Technology). During the tests, the buffer output at the excitation frequency of 20 kHz was continuously measured using a 16-bit digitizer (NI myDAQ; National instruments) with 200 kHz sampling rate. A custom Labview program (Labview 2018; National Instruments) was developed to detect and record the peak-to-peak amplitude of the buffer output that is directly proportional to the luminesce intensity of Ru-dye immobilized in the O_2_-sensing film. The collected data were averaged every 3 and 12 hours in Fig. 2f and 2g, respectively.

### Design of the external transceiver

The external transceiver consisted of transmitter (TX) and receiver (RX) paths. The TX path included a commercial high-voltage pulser with an integrated TX/RX switch (MAX14808; Maxim Integrated) and a digital controller module (NI PXIe-6363; National Instruments). During the TX mode, the high-voltage pulser converted a low-voltage signal from the digital controller module to a high-voltage signal, necessary to drive an external ultrasound transducer to generate ultrasound pulses. The RX path included an ultralow noise amplifier (AD8432; Analog Devices) to receive and amplify the backscatter signal from the external transducer, a gain amplifier to further amplify the signal to a level within the input range the analog-digital converter (ADC), and a digitizer with an antialiasing filter and a 14-bit high-speed ADC (NI PXIe-5122; National Instruments) to filter and digitize the signal after receiving and amplification. In addition to the switch integrated into the pulser, an extra digitally-controlled switch (ADG619; Analog Devices) was used to minimize the electrical coupling (interaction) between the TX and RX paths. The TX and RX paths were synchronized to each other by using the same reference clock integrated into the backplane of the PXI chassis (NI PXIe-1062Q; National Instruments). Note that the digital controller, digitizer, and NI PXIe-8360 modules were inserted in the chassis, in which the NI PXIe-8360 module was used to connect the chassis to a computer for communication with the other modules and data transfer.

A custom Labview program (Labview 2018; National Instruments) was developed to control the modules and to process the backscatter data in real-time. A (TX and RX) communication protocol was encoded in the program. During real-time data processing, the backscatter data digitized by a 14-bit ADC with a sampling rate of 20 MHz were resampled by a factor of five and then interpolated with a sinc function. The sinc interpolation was followed by a peak detection to extract the envelope of the backscatter signal and linear interpolation to increase the number of data points and hence to improve the accuracy in the determination of an optimal threshold value that minimizes bit error rate (BER). An optimal threshold (that is, the half value of the sum of modulated and unmodulated backscatter signal amplitudes) was determined by taking the mean of the data points from the time intervals where the steady-state backscatter signal was amplitude modulated and unmodulated. The threshold was used to convert the digitized data into digital format: bits (“0” or “1”). The bits were scanned to find a preamble and a postamble and hence to extract data bits. The binary coded data (bits) were converted to numeric data, which was stored on a computer.

### *In vitro* and *ex vivo* characterization

An *in vitro* characterization of the wireless oxygen monitoring system was performed in a custom-built water tank using a 25.4 mm diameter, 2.25 MHz single-element external ultrasonic transducer (V304-SU-F1.88IN-PTF; Olympus) with a focal depth of 47.8 mm, mounted on manual translation stages (Thorlabs) and connected to an external transceiver board, at various alignments and positions of the wireless oxygen sensor with test leads, mounted on top of a steel rod with a diameter of 0.75 mm connected to a manual rotation stage (Thorlabs). In measurements, the external transducer face was covered with a thin sheet of latex by filling the empty space between the transducer face and the latex sheet with castor oil (used as a coupling medium), to protect the matching layer of the transducer from possible damage due to the long-time direct contact with water or ultrasound gel. A hydrophone (HGL-0400; Onda) was used to calibrate the output pressure and hence the acoustic intensity and to characterize the acoustic beam patterns of the external transducer (Supplementary Figs. 8).

In measurements, the water tank was placed on a stirring hotplate (Thermo Scientific Cimarec), to keep the water temperature constant at 37 ± 0.1 °C, to simulate physiological temperature, and to stir using a magnetic stirrer to increase the speed of a transition from low to high O_2_ level and vice versa in distilled water. Water O_2_ concentration was monitored using a commercial O_2_ probe with a 300 µm core diameter (NEOFOX-KIT-PROBE; BIFBORO-300-2; Ocean Optics) varied by controlling the ratio of O_2_ and N_2_, supplied to the water tank through two pipes, via a matched pair of gas flow controllers (FMA-A2407; Omega) connected to O_2_ and N_2_ gas cylinders. A customized Matlab program controlled the gas flow controllers through a digital-to-analog converter board (NI myDAQ; National instruments) connected to a computer.

The measured phase output data from the wireless system were converted to O_2_ concentration (partial pressure of oxygen, pO_2_) in mmHg by an exponential equation: pO_2_ (mmHg) = A·e^(B/Ф)^ + C, where Ф is the phase output, and A, B and C are constant coefficients obtained from curve fitting (see Supplementary Fig. 6).

Ethylene oxide (EtO) sterilization with an exposure time of 4 h at 37 ± 3 °C and an aeration time of 24 h at 37 ± 3 °C was performed by a commercial vendor (Blue Line Sterilization Services LLC, Novato, CA).

To assess the functionality of the fully-packaged O_2_ sensors (Fig. 1f) over time in an environment that mimics (to first order) in vivo biofouling, phosphate-buffered saline (PBS, 1×) without calcium and magnesium (Corning; Mediatech Inc.) and pooled human serum (off the clot) (purchased from Innovative Research, Inc., Novi, MI) were used for 10-day incubation of the sensors at 37 ± 0.1 °C in an oven (Test Equity Model 107). The two antibiotics, penicillin and streptomycin with a final concentration of 100 units/mL and 100 μg/mL (Gibco by Life Technologies, Catalog # 15-140-122; purchased from ThermoFisher Scientific), were added to human serum to inhibit bacterial growth during the study. The test in serum was performed by placing the sensor in a container, where antibiotics-added serum was replaced every 24 h to ensure sterile conditions during the length of the study. In the fouling tests, the sensors were operated at 350 samples per second (Hz) sampling rate with differential, 2 MHz AC signals produced by a Keysight 33500B function generator, which are ac-coupled to the rectifier inputs of the IC. One of the rectifier inputs was connected to a high-input impedance buffer amplifier (LTC6268; Linear Technology). The buffer output, O_2_ data, was recorded by a 14-bit high-speed digitizer (NI PXIe-5122; National Instruments), synchronized to the clock of the function generator, and a custom Labview program (Labview 2018; National Instruments).

Measurements for uplink performance assessment of the system at 10 cm depth in DI water and through a fresh ex vivo porcine specimen were performed with a custom-designed and -built spherically-focused, 2 MHz, 25.4 mm diameter ultrasonic transducer with a focal length of 88.1 mm (Sensor Networks Inc.) (see Supplementary Fig. 9 for the transducer beam pattern). In the *ex vivo* measurements, a porcine tissue sample was positioned between the wireless sensor and the external transducer, with coupling enabled by ultrasound gel (Aquasonic Clear; Parker Labs). Air bubbles in ultrasound gel were removed via centrifugation at 2800 rpm for 10 min. In order to remove possible air bubbles entrapped between the sensor and the tissue, the sensor was positioned on a tissue sample in a container filled with DI water. A piece of ultrasound absorbing material was placed under the tissue sample to avoid ultrasound reflection from the bottom interface of the container.

### Backscatter modulation depth

Backscatter relative difference is defined as the ratio of the amplitude difference between the modulated and unmodulated backscatter signals to the amplitude of the unmodulated backscatter signal. The modulation depth percentage was calculated by multiplying the backscatter relative difference by 100. Backscatter relative difference plots were obtained by collecting data samples from the time points where the steady-state backscatter signal was amplitude modulated and unmodulated during the O_2_ measurement.

### Ultrasound (US) link power transfer efficiency

The US link power transfer efficiency is defined as the ratio of the electrical input power of the IC to the acoustic power emitted from the external transducer, which depends on the beam focusing ability of the external transducer, the frequency-dependent attenuation of US intensity in the propagation media, and the power conversion efficiency of the sensor. The acoustic power at the transducer surface was calculated by integrating the acoustic field intensity data, obtained by a hydrophone at the focal length, over a circular area where the intensity of the side lobes is not negligible. The power conversion efficiency of the sensor, relying on the receive (acoustic-to-electrical conversion) efficiency of the piezo and the impedance matching between the piezo and the IC, is equal to the ratio of the electrical input power of the IC to the acoustic power at the surface of the sensor piezo; the acoustic power at the piezo surface was calculated by integrating the acoustic field intensity data from the hydrophone over the surface of the sensor piezo.

### *In vivo* measurements

The Committee on Animal Research of the University of California, San Francisco, approved all protocols and procedures. All animal research conforms to the National Institutes of Health (NIH) guidelines as outlined in the Guide for the Care and Use of Laboratory Animals.

“Lambs (2-12 weeks of age) are given ketamine (10-15 mg/kg, IM). An appropriatly sized catheter is placed percutanelouly into a peripheral vein. The lambs are given intravenous (IV) boluses of fentanyl citrate and diazepam for sedation, and then anesthetized with continuous intravenous infusions of fentanyl citrate, ketamine hydrochloride and diazepam. Heart rate and blood pressure are continuously monitored, and doses adjusted with changes in these parameters. Then catheters are placed in an artery and vein of each hind limb. This procedure is done either before or after endotracheal intubation, when a surgical plane of anesthesia is present. Separate arterial lines are sometimes needed such that arterial blood sampling, for arterial blood gas management, as well as for experimental analysis is possible without interrupting continuous hemodynamic monitoring. This is particularly important to monitor anesthesia and the health of the lamb during the study. Likewise, separate venous access is needed to allow for the continuous administration of maintenance fluids and anesthesia while simultaneously allowing intravenous access for the administration of bolus fluids, blood products, experimental agents, and/or other intravenous agents as needed throughout the study. The lambs are given IV boluses of fentanyl citrate and diazepam and then intubated with a 4.5-7.5 mm OD cuffed endotracheal tube and mechanically ventilated with a pediatric time-cycled, pressure-limited ventilator. Vecuronium bromide is given intermittently (IV bolus) for muscle relaxation, and redosed for movement. Ventilation with 21% oxygen is adjusted to maintain a systemic arterial pCO2 between 35 and 45 torr. To ensure lambs are adequately anesthetized during paralysis, heart rate, systemic arterial pressure, temperature and response to stimuli are continuously monitored. In addition, arterial blood gas readings are performed every 30-120 mins to determine pCO2, pO2, and pH parameters. In general, blood gases will be obtained q 30 min. However, if blood gases are stable for several hours, then blood gas sampling will be expanded to a maximum of 2 hours to decrease blood withdrawal. The anesthetic level will not be decreased after the vecuronium has been administered; however, it can be increased if clinical signs of an inadequate level of anesthesia are observed.”

In this study, juvenile or adult lambs (n = 2 animals (A and B)) were anesthetized with fentanyl, ketamine, and diazepam and paralyzed with vecuronium to facilitate intubation and mechanical ventilation. Ongoing sedation and neuromuscular blockade were administered as a continuous infusion of ketamine, fentanyl, diazepam, and vecuronium. The anesthesia mixture was titrated to maintain age-appropriate HR. Femoral venous and arterial access were obtained via cutdown of the hind limbs, and arterial pressure was continuously transduced and recorded. The animals were ventilated with a positive end expiratory pressure of 5cm H_2_O, tidal volumes of 10mL/kg, and respiratory rate titrated to maintain pCO_2_ of 35-45 millimeters mercury (mmHg) by arterial blood gas measurements. An oxygen blender allowed rapid titration of fraction of inspired oxygen (FiO_2_). Following instrumentation, the animals were allowed to recover to steady state until they required no further adjustment to sedatives and exhibited stable hemodynamic parameters. This time was designated as the normoxic baseline and blood gas analysis was performed to confirm physiologic parameters.

In order to facilitate O_2_ sensor placement, a 2-way skin flap of appropriate size was created over the lateral thigh. The uppermost layers of skin were carefully dissected away from the subcutaneous space to ensure tissue integrity and then reflected away from the site. Exposure of the posterior fascial compartment was then visualized and dissected away to reveal the biceps femoris muscle. Because of muscle depth, the lateral aspect of the biceps femoris was carefully dissected to create a tunnel into the underlying fascial plane. The wireless O_2_ sensor was then seated under the biceps femoris via the created tunnel and skin closed in a simple continuous fashion. Note that the wireless O_2_ sensor had five flexible leads attached to it only for electrical testing and debugging to make sure that the sensor operates properly before *in vivo* measurements, and, as seen in Fig. 5a-d, they were not even accessible from outside except two leads (ground and V_DC-DC_) in the *in vivo* measurements, which were only used to check the sensor functionality after implantation and completely disconnected during measurements. A commercial wired O_2_ probe (NEOFOX-KIT-PROBE; BIFBORO-300-2; Ocean Optics) was placed nearby to confirm tissue pO_2_ measurements. Tissue pO_2_ measurements were compared with arterial pO_2_ (paO_2_) obtained by blood gas analysis, as well as with blood hemoglobin O_2_ saturation (SpO_2_) obtained via a GE Dash 3000 patient monitor and GE Masimo finger probe placed on the ear.

At the end of the protocol, all lambs were euthanized with a lethal injection of sodium pentobarbital (150 mg/kg) followed by bilateral thoracotomy as described in the National Institutes of Health Guidelines for the Care and Use of Laboratory Animals.

*In vivo* experiments were carried out using a 12.7 mm diameter, 2.25 MHz single-element external ultrasound transducer with a focal depth of 21.6 mm (V306-SU-F0.88IN-PTF; Olympus), demonstrating a similar beam pattern with the 25.4 mm diameter transducer used in the *in vitro* experiments. The output pressure and acoustic intensity for the 12.7 mm diameter transducer was characterized in a water tank using a hydrophone (HGL-0400; Onda) after its face was covered with a thin sheet of latex and castor oil, to protect its matching layer from possible damage due to a long-time direct contact with ultrasound gel. The external transducer was attached to a custom five-axis micromanipulator, built using single-axis linear stages (DT-100 X; SD Instruments), a tilting stage (MM-2A; Newport), and a 0.5 mm acrylic sheet. The micromanipulator stage, used for alignment of the external transducer to the implanted sensor, was placed on top of the skin layer on a 7 mm-thick acoustic standoff pad (DIY ultrasound phantom gel; Humimic Medical) with ultrasound gel (Aquasonic Clear; Parker Labs) applied to the skin surface and the external transducer for acoustic coupling between the tissue and the transducer. The ultrasound backscatter amplitude was used for fine alignment of the implant and the external transducer.

Tissue pO_2_ measurements were performed with the wireless system operated at a sampling rate of 350 samples per second. In the *in vivo* measurements, the maximum distance from the external transducer to the wireless O_2_ sensor, operated with an acoustic field that had a derated I_SPTA_ of 454 mW/cm^2^ and a mechanical index of 0.08 (both below the FDA regulatory limits of 720 mW/cm^2^ and 0.19), was ∼26 mm with ∼19 mm consisting of tissue (including skin, fat, and muscle). The distance between the implanted sensor and the external transducer was estimated from the round-trip time-of-flight (that is, the time delay between the received backscatter signal from the sensor piezo and the signal that drove the external transducer). Both the wireless and the wired data, shown in Fig. 5, were averaged every 5 s. Two identical wireless O_2_ sensors were used in *in vivo* measurements; the first sensor response to various O_2_ concentrations in water and animal A was shown in Figs. 4e and 5e, and the second sensor response in water and animal B was shown in Supplementary Fig. 10 and Fig. 5f. All images were captured by a smartphone camera.

### Data availability

All data supporting the results in this study are available within the article or its Supplementary Information.

### Code availability

The custom Labview program and Matlab codes used in this study are available from the corresponding author upon reasonable request.

## Acknowledgements

This work was supported by the Chan Zuckerberg Biohub (CZB) (to M.M.M.) and by NIH/NICHD R44HD094414, R01HD072455 (to J.R.F. and E.M.). The authors would like to thank Courtney Losser, Rachel Hutchings and Christian Vento for expert assistance with animal handling and surgery, as well as members of the Laboratory Animal Resource Center (LARC) at University of California, San Francisco. The authors would also like to thank the Berkeley Wireless Research Center and Professor Rikky Muller (University of California, Berkeley) for access to IC design software.

## Author contributions

S.S. supervised the project, designed and built the wireless system, designed and performed the *in vitro* and *ex vivo* experiments, and analysed and interpreted the associated data. S.S., J.R.F. and E.M designed and performed the *in vivo* experiments and interpreted biological data. M.M.M contributed to the design of the experiments. S.S., E.M. and M.M.M. participated in writing the paper. All authors contributed to the discussion of the paper.

---

**Supplementary Fig. 1.**
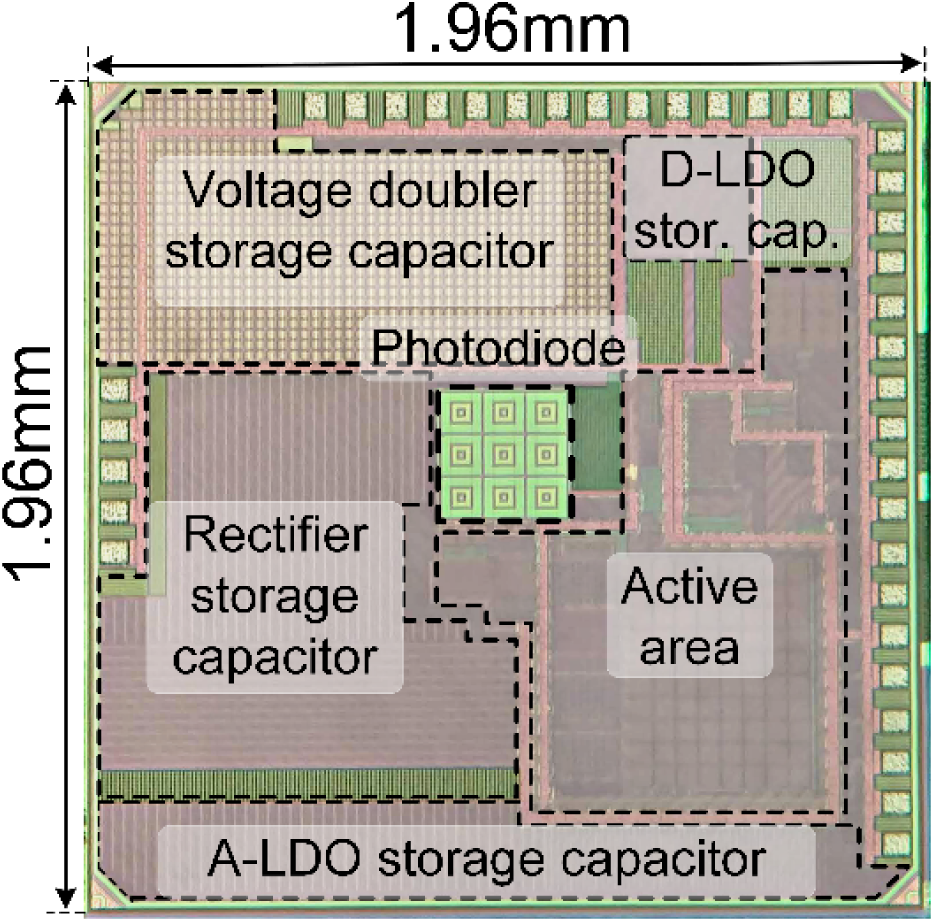
Sensor IC die micrograph.

**Supplementary Fig. 2.**
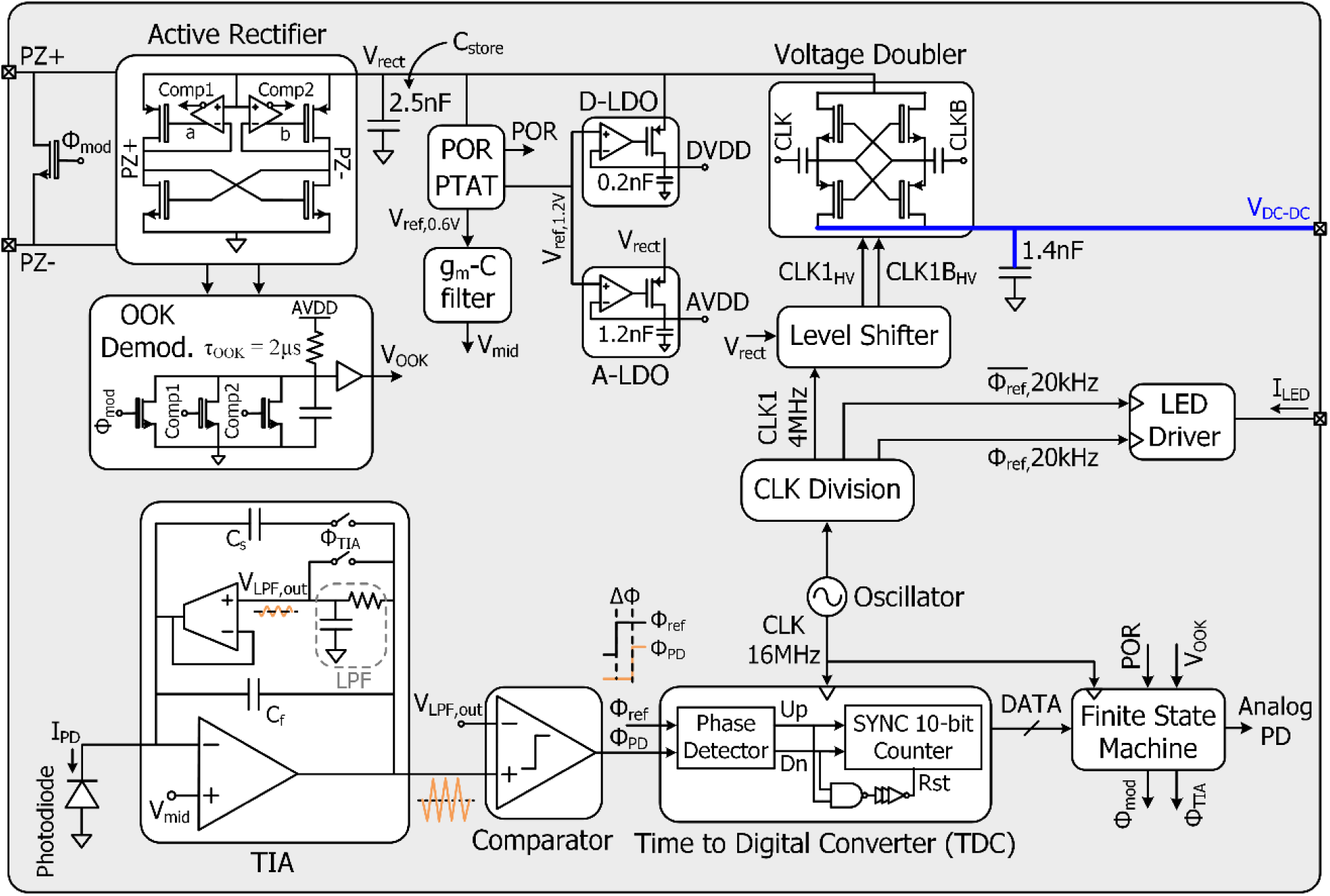
A more detailed schematic of the IC architecture. An analog front-end (AFE) consisted of a transcapacitance amplifier, in which the DC feedback was provided using an active biasing circuit and the switches, controlled by Ф_TIA_, were implemented to minimize the settling time after duty-cycling, and of a comparator. The rectifier comparator outputs (Comp1 and Comp2) and the modulation signal (Φ_mod_) were used as inputs to the OOK demodulator that detected the envelope of the downlink signal and generated a notch. The TDC was based on a 10-bit synchronous counter and a phase detector. An LED driver was implemented using an 8-bit current digital-to-analog converter (DAC) that was driven by the 20 kHz complementary clock signals to avoid the rectifier voltage (V_rect_) fluctuation. A cross-coupled voltage doubler was designed to boost V_rect_.

**Supplementary Fig. 3.**
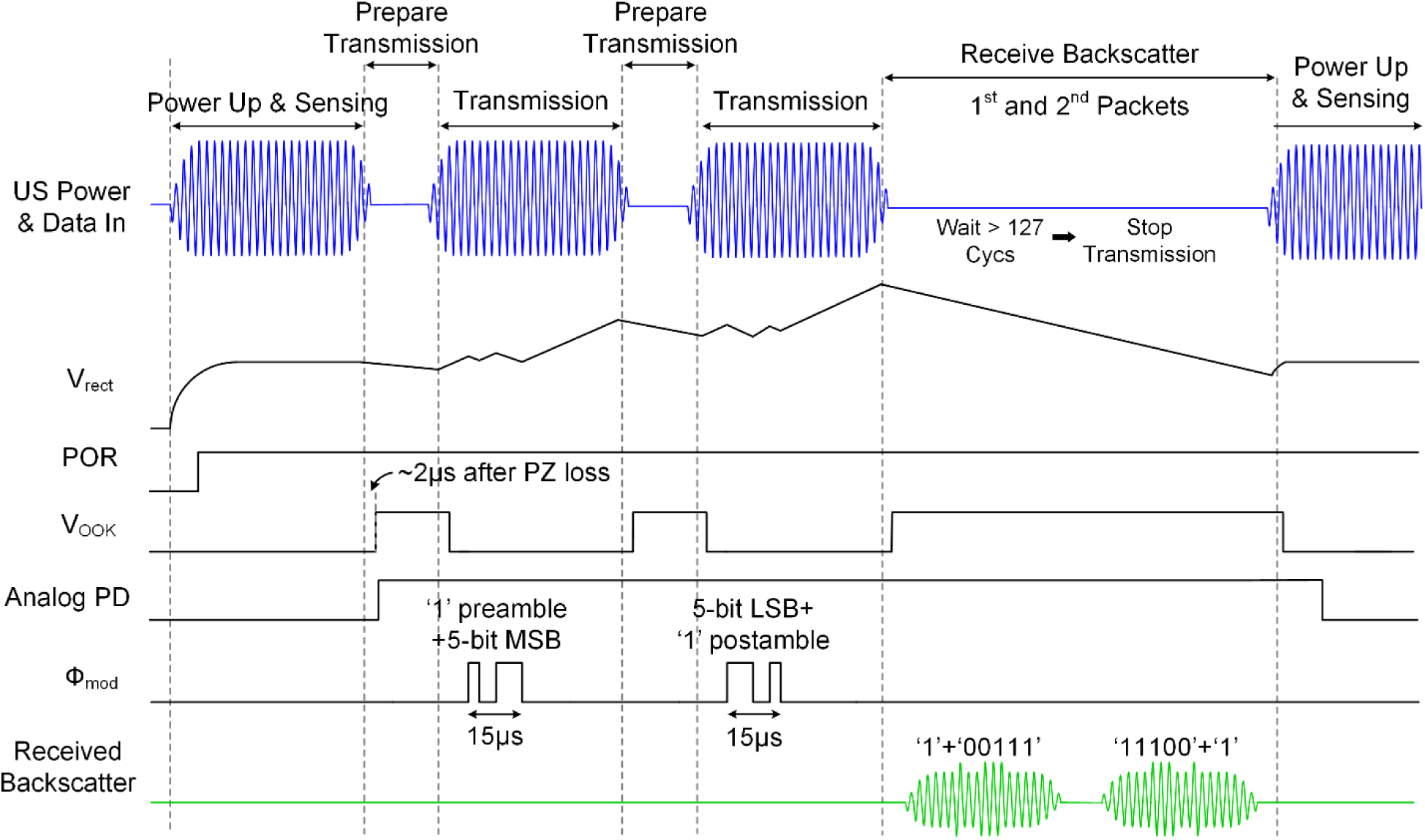
An alternative timing diagram for a wireless O_2_ sensor operated at a depth of greater than 5 cm. Here, the two data packets were subsequently transmitted, as in Fig. 3b, but received in a single ultrasound-free interval. As an alternative to the communication protocol demonstrated in Fig. 3b, this protocol enables to reduce the time spent during data transmission, which in turn allows increasing the sampling rate of the system.

**Supplementary Fig. 4.**
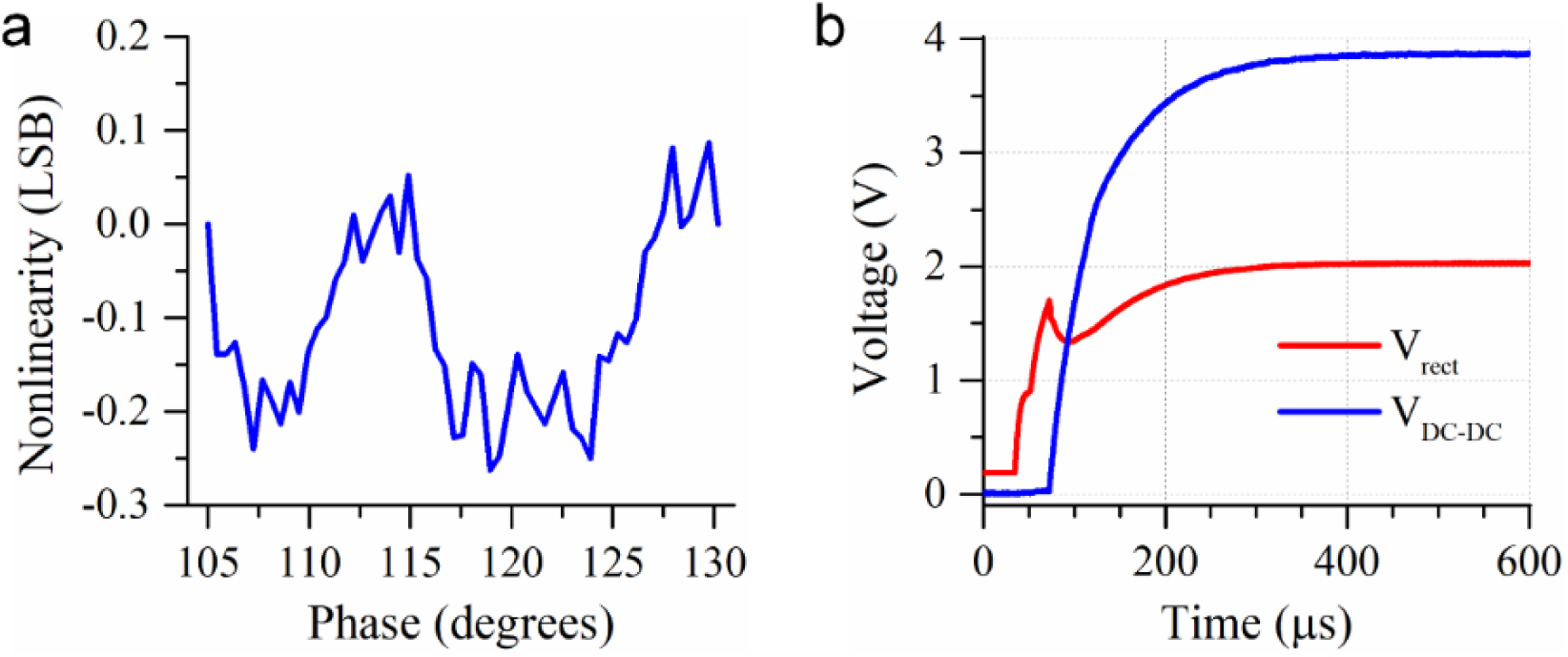
Measured waveforms of the IC. **a**, Nonlinearity in the phase readout circuitry. **b**, A rectifier voltage (V_rect_) and a voltage doubler output (V_DC-DC_) during a power-up period of ∼150 µs. At steady- state, the voltage conversion ratio (VCR) of the voltage doubler was 1.91.

**Supplementary Fig. 5.**
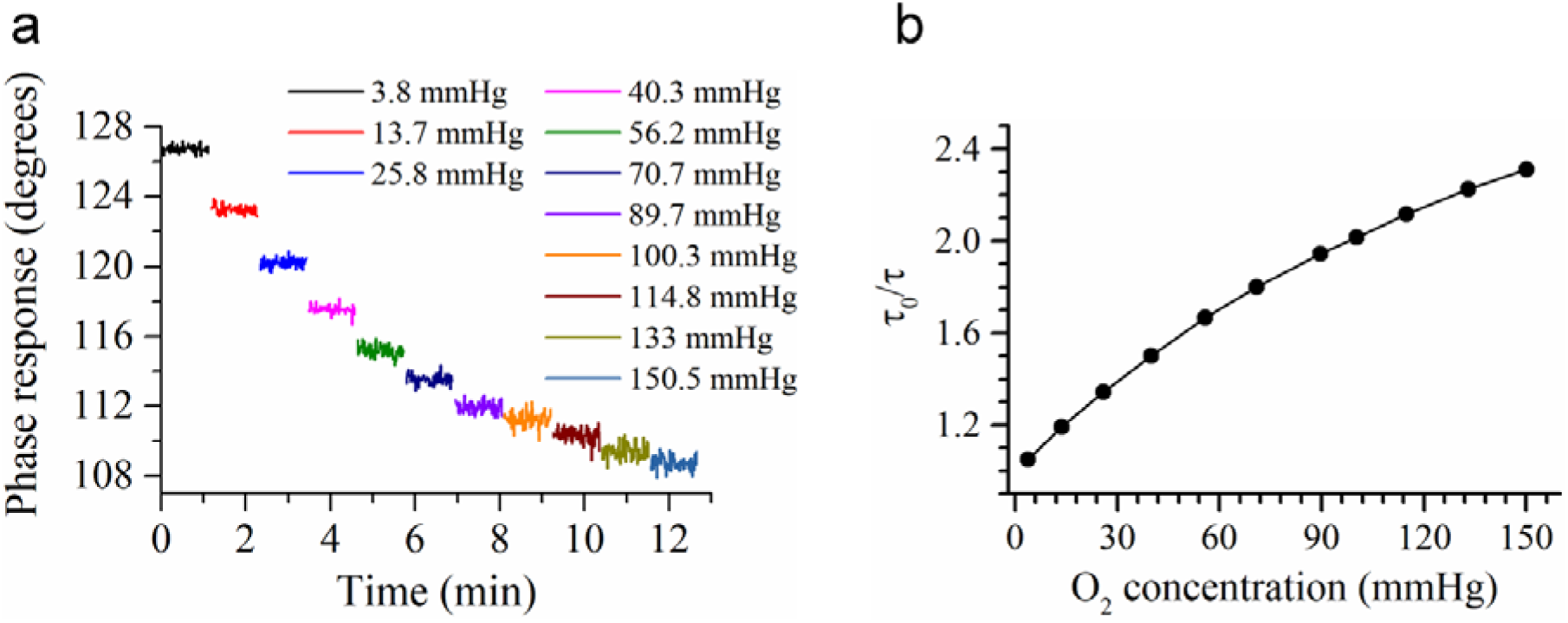
Wireless system response to various O_2_ concentrations in the range of 3.8-150 mmHg. **a,** The measurement was recorded with a wireless O_2_ sensor operated at 350 samples per second sampling rate and 5 cm depth during the *in vitro* characterization (see Fig. 4). **b,** Lifetime (τ)-based Stern-Volmer plot obtained from the data shown in (**a**).

**Supplementary Fig. 6.**
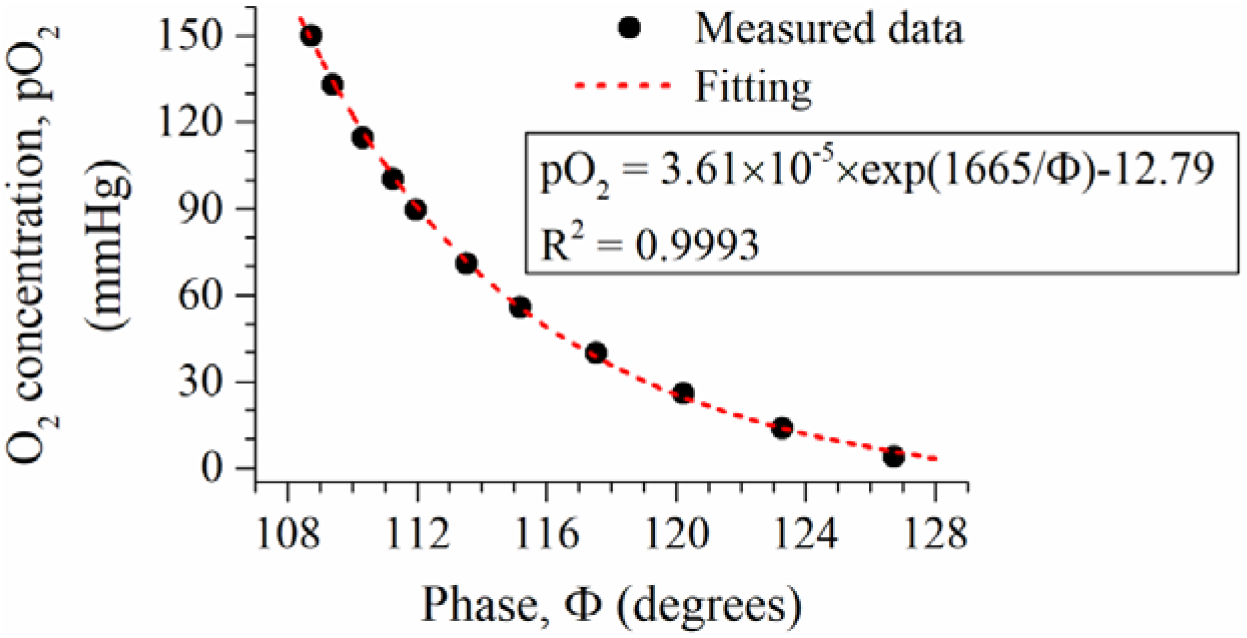
Calibration curve of the wireless O_2_ sensor. Calibration of the O_2_ sensor in distilled water at 37 °C was conducted by periodically increasing the O_2_ concentration of water surrounding the sensor, as shown in Supplementary Fig. 5a. The dissolved O_2_ concentration was measured at each step by a commercial O_2_ sensor (NEOFOX). The calibration curve was fitted by an exponential equation: pO_2_ (mmHg) = A·e^(B/Ф)^ + C, where Ф is the sensor phase output, and A, B and C are constant coefficients obtained from the curve fitting. The exponential equation provides high accuracy with an R^2^ of 0.9993. Alternative equations, such as high-order polynomial functions, could be used for calibration curve fitting.

**Supplementary Fig. 7.**
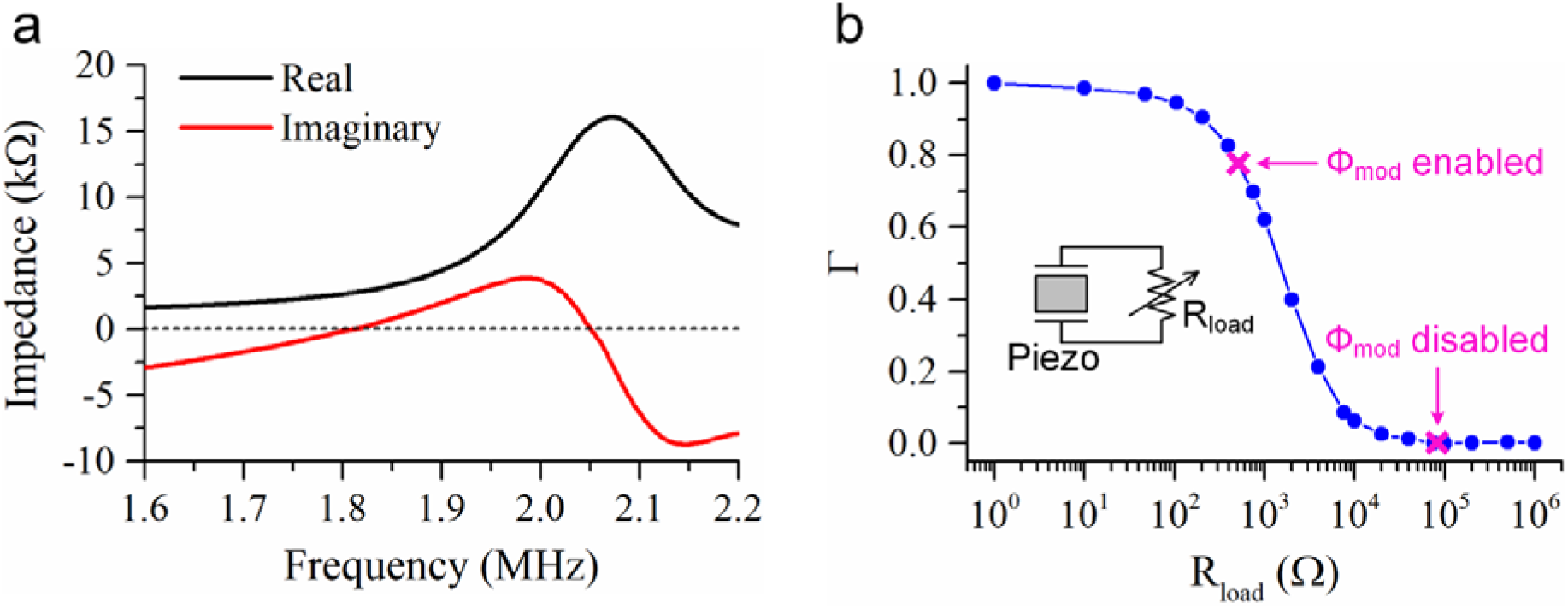
Impedance and backscatter characterization of a 750 µm × 750 µm ×750 µm PZT piezoelectric crystal. **a,** Measured impedance of the 10 μm-thick parylene-coated piezo crystal as a function of frequency in distilled water. During system operation, the piezo was driven at 2 MHz frequency that is close to its open-circuit resonance frequency of 2.05 MHz. The impedance values at 2 MHz provide a good impedance matching with the rectifier input resistance (R_in_) of ∼11.8 kΩ at the desired (2 V) output voltage of the rectifier, yielding an impedance matching efficiency of ∼97% between the piezo and the rectifier. A capacitive matching network^87^ could be used to improve matching efficiency further. **b,** Normalized acoustic reflection coefficient (Г) of the piezo crystal versus load resistance (R_load_), measured with ultrasound at 2 MHz. Here R_load_ simulates R_in_.

**Supplementary Fig. 8.**
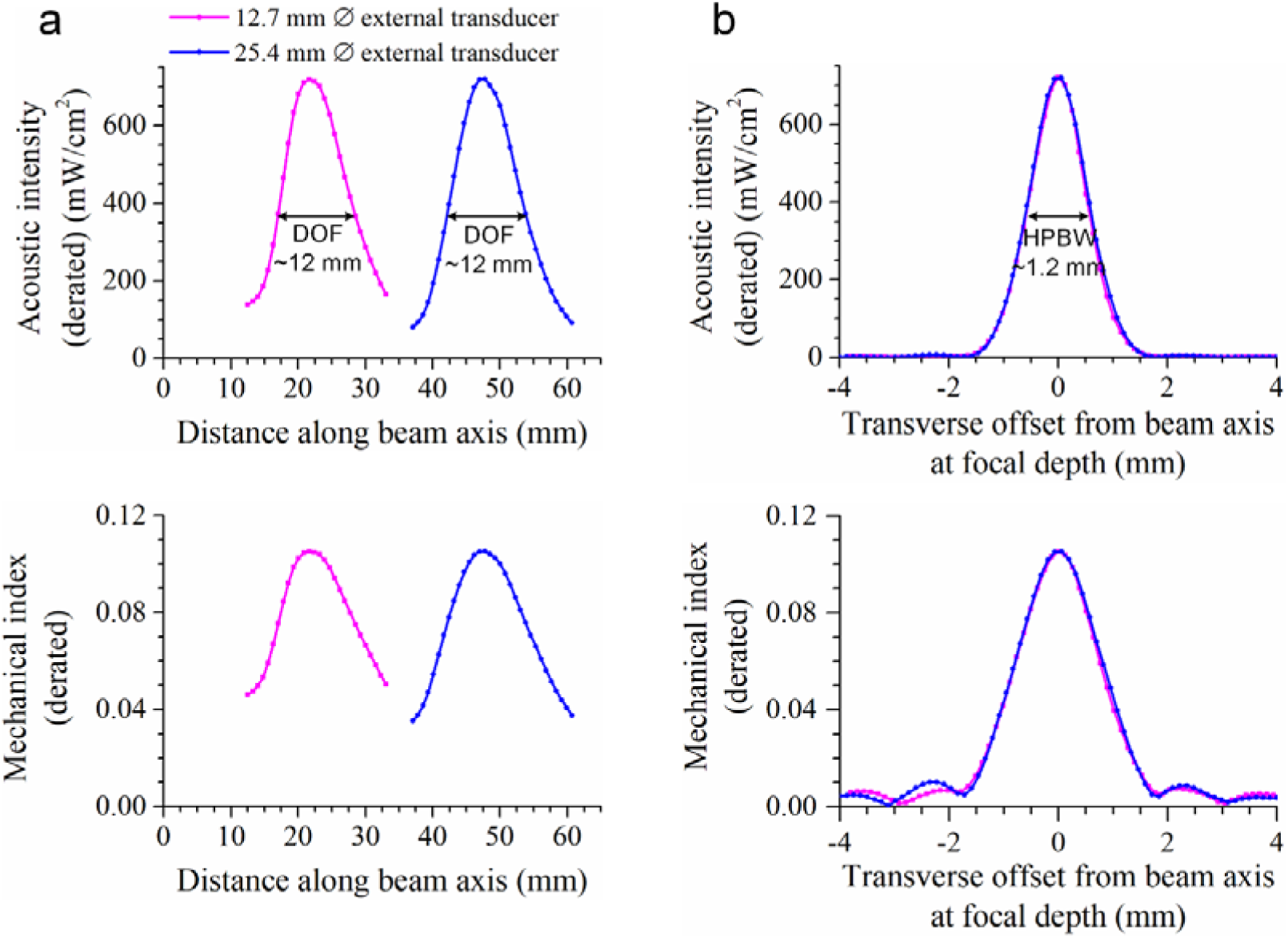
Characterization of the external ultrasound transducers, used for measurements at moderate depths ≤5 cm in distilled water using a hydrophone. **a,** Longitudinal beam patterns. **b,** Transverse beam patterns.

**Supplementary Fig. 9.**
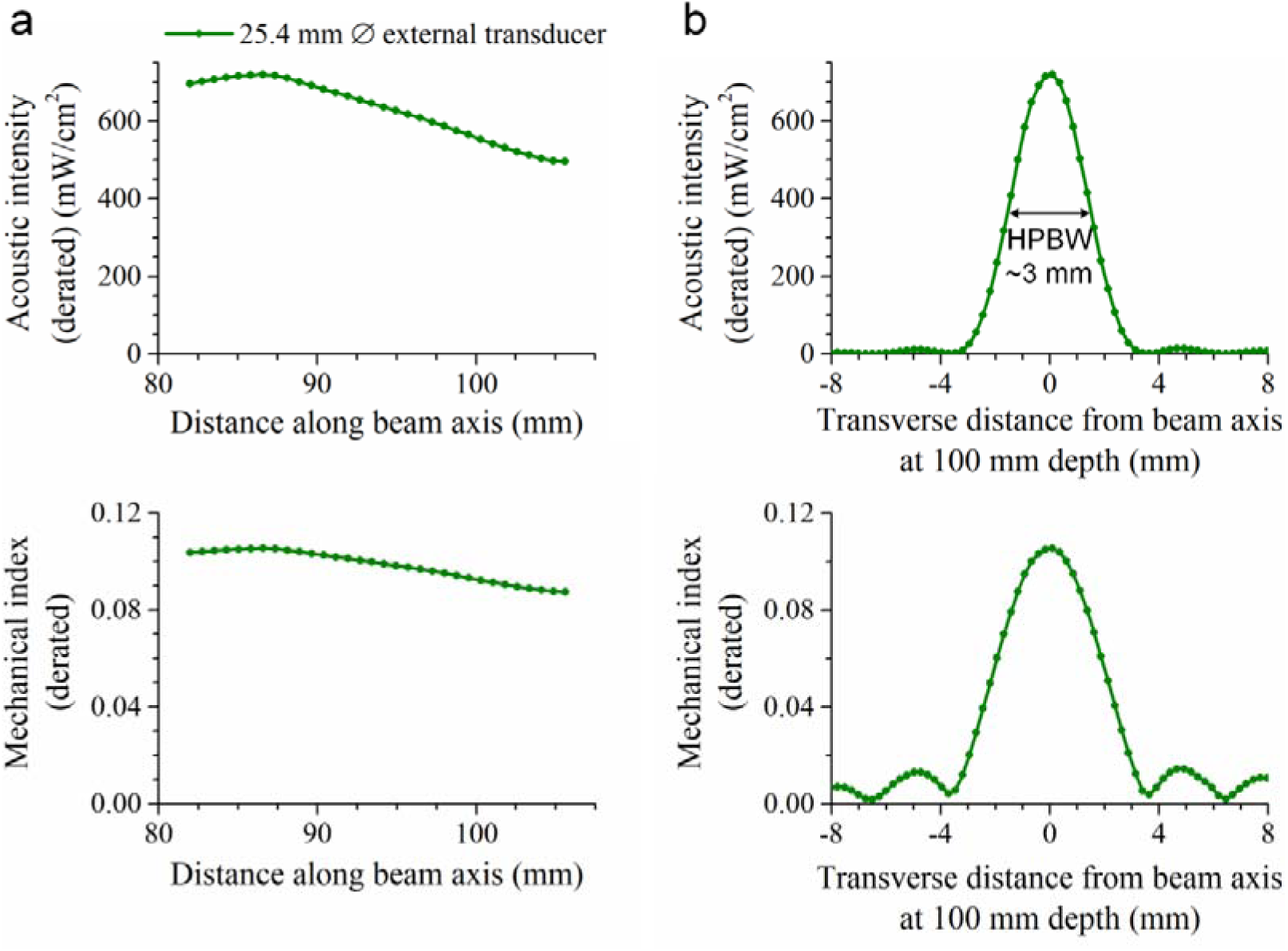
Characterization of the external ultrasound transducer, used for measurements at 10 cm depth in distilled water using a hydrophone. **a**, Longitudinal beam pattern. **b**, Transverse beam pattern.

**Supplementary Fig. 10.**
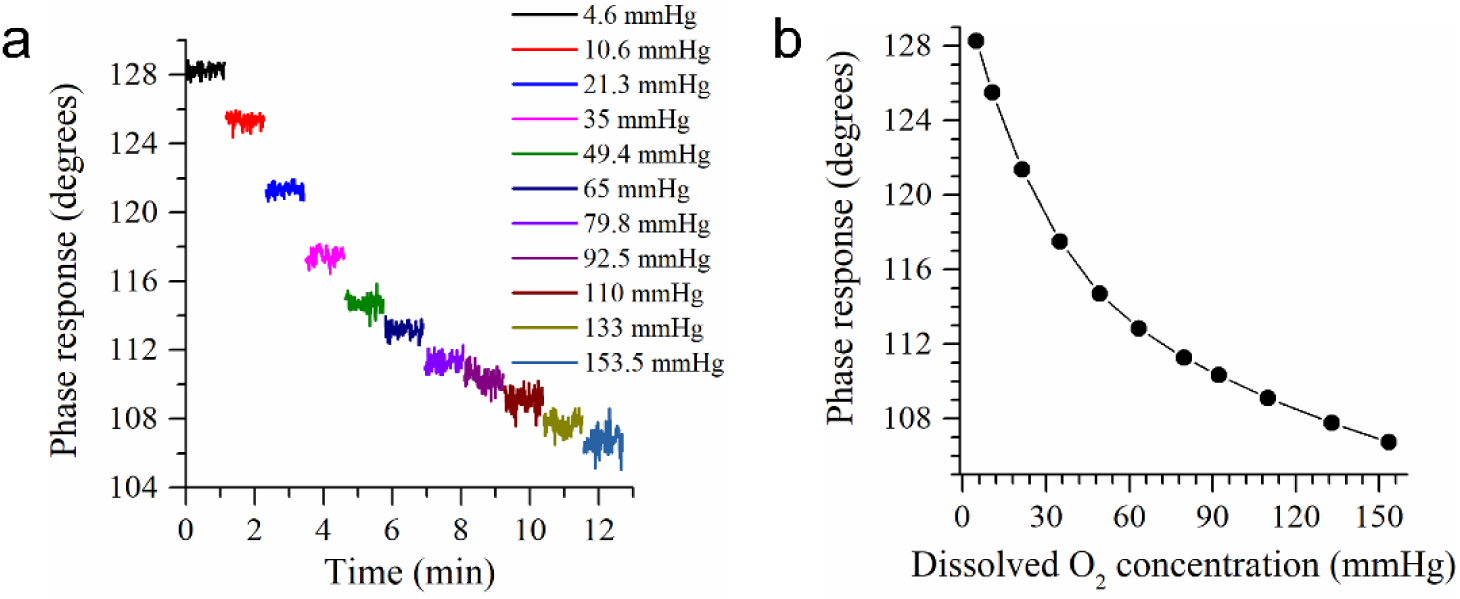
System response to various O_2_ concentrations. An additional, identical wireless O_2_ sensor was also characterized in distilled water at a depth of 5 cm and a sampling rate of 350 samples per second at various O_2_ concentrations. This sensor was also used for tissue O_2_ monitoring. The data, shown in Fig. 5f, were collected using this sensor. **Effect of ultrasound (US) link misalignment on the system operation.** Although the use of acoustic waves, instead of near-field electromagnetic waves^18^, enabled to power and communicate with the mm-scale wireless O sensor at great depths (≥5 cm), it made the system sensitive to the US link alignment between the external transceiver and the wireless O_2_ sensor. Therefore, it was crucial to understanding the impact of US link misalignment on the system operation. The misalignment sensitivity of the system was evaluated by measuring the sensor V_rect_ and the uplink bit error rate (BER) (Supplementary Fig. 11). The minimum V_rect_ necessary to turn on the µLED and hence to operate the sensor was ∼1.36 V (see Supplementary Fig. 12a). The minimum acoustic intensity required to produce 1.36 V V_rect_ was 142 mW/cm^2^, ∼19.7% of the FDA limit (derated I_SPTA_) of 720 mW/cm^2^. This ∼5× acoustic intensity margin provided an ability to tolerate US link misalignment for proper sensor operation while keeping the intensity at safe levels. Alignment measurements were performed in a water tank using a 25.4 mm-diameter transducer with a focal depth of 47.8 mm, generating acoustic pulses at 2 MHz with a fixed I_SPTA_ of 220 mW/cm^2^ (Supplementary Fig. 11a). When the sensor was operated on the center axis of the US beam with a zero angular offset, the system could robustly operate in a wide depth range of approximately 41-57 mm without sacrificing the BER performance (Supplementary Fig. 11b); measured BERs at various depths within the operating depth range were below 10^−5^. When the sensor, positioned on the central axis of the acoustic field at the focal depth, was scanned transversely and angularly relative to the central axis of the sensor’s piezo, the system also functioned properly with a 24° angular and 0.56 mm transverse offset relative to the beam central axis, at the expense of slight BER performance degradation at different angular and/or transverse offsets (Supplementary Fig. 11c,d). The link misalignment measurements showed that the system operation exhibited relatively high tolerance to the depth misalignment compared to the transverse misalignment, as the -3 dB depth-of-field (DOF: ∼12 mm) of the single-element focused transducer is substantially higher than its beam spot size (HPBW: ∼1.2 mm) at the focal depth (Supplementary Fig. 8). In practice, a fine depth alignment could be achieved by using either an acoustic standoff pad and/or ultrasound gel or an external transducer with differing focal depths. Angular alignment within the operating angular misalignment range of ±24° could be performed by careful surgical placement of the sensor in tissue. Here operation was relatively sensitive to transverse misalignment due to the transverse beam pattern produced by the external transducer. The transverse beam spot size at the desired focal length could be increased by optimizing the geometry of the acoustic lens that was built into the transducer^89, 90^. A custom-designed and -built single-element transducer with a wider spot size could be used to reduce the system sensitivity to transverse misalignment, but at the expense of reduced US link power transfer efficiency.

**Supplementary Fig. 11.**
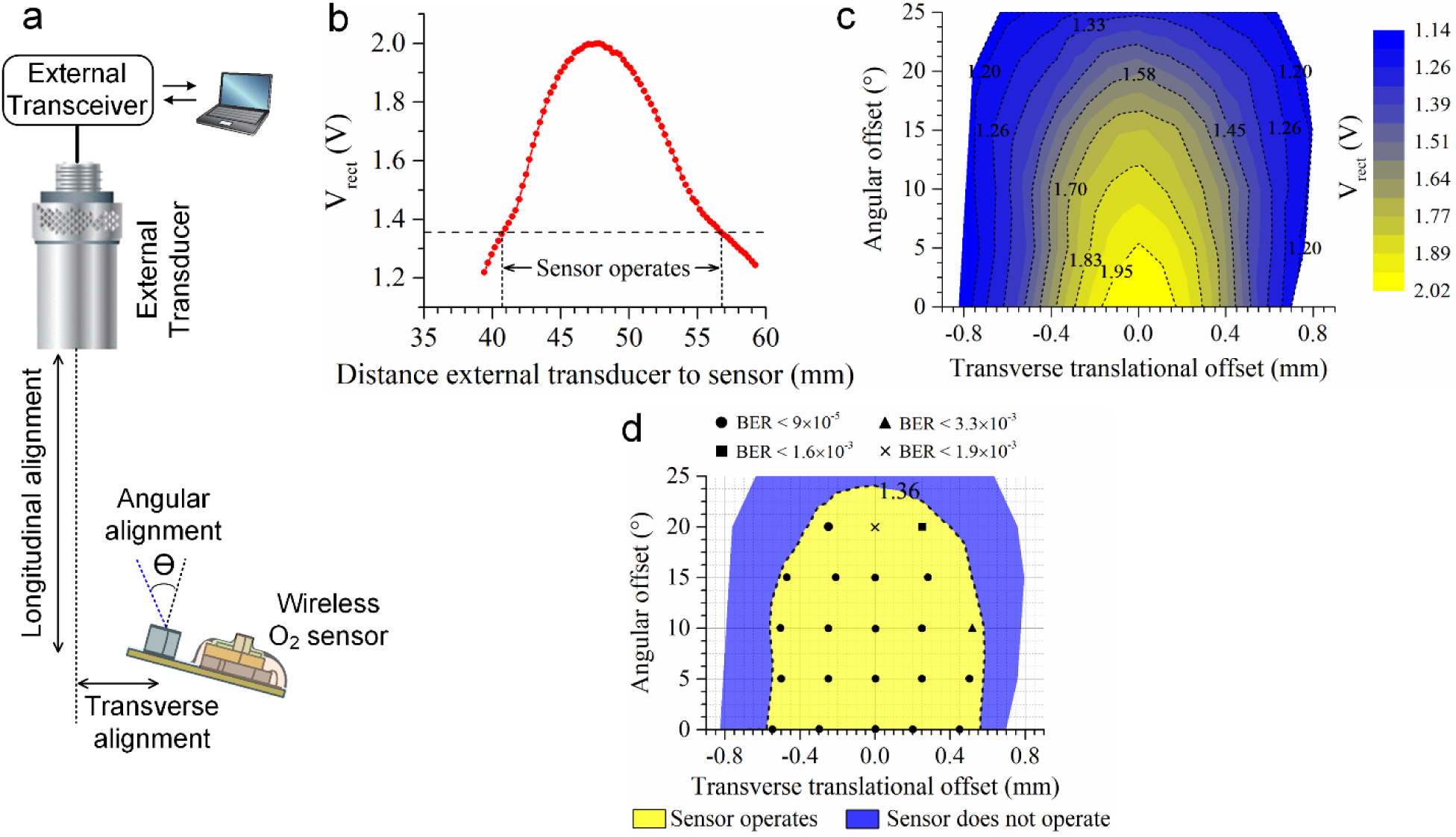
Effect of ultrasound (US) link alignment on the system operation. **a**, Schematic diagram of the misalignment parameters. The measurements were performed in distilled water using a spherically-focused external transducer with a 25.4 mm diameter and a focal depth of 47.8 mm. **b**-**d**, A wireless sensor was operated at a fixed I_SPTA_ of 220 mW/cm^2^. **b**, A sensor was operated while its depth was longitudinally scanned along the central axis of the acoustic field, showing a wide operating window of 16 mm. **c**, A wireless sensor was placed aligned to the center of the focal plane at the focal depth, and its position and orientation were scanned along transverse and angular directions relative to the central axis of the sensor piezo. **d**, Color map depicted a region where the sensor operates.

**Supplementary Fig. 12.**
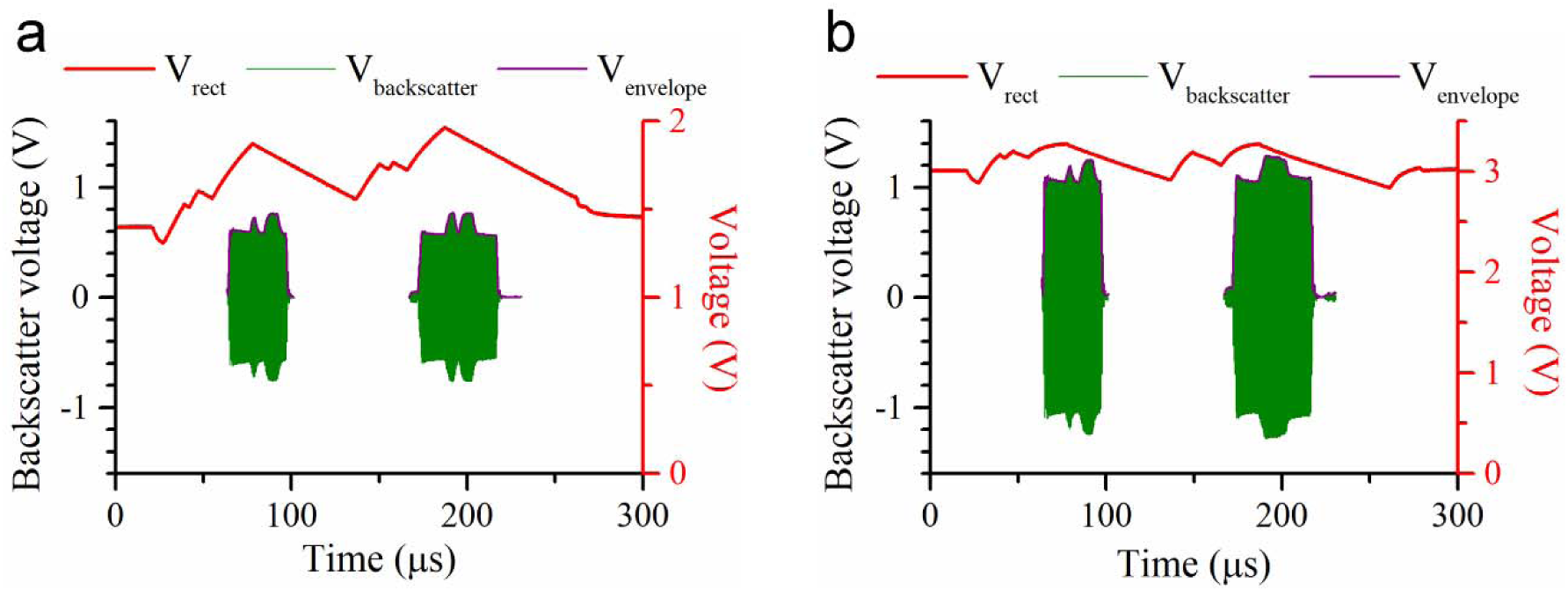
Wireless measurement of a single O_2_ sample for the sensor operated at 5 cm depth in water with different acoustic intensities. **a, b**, The wireless O_2_ sensor was operated with a 2 MHz acoustic wave with an I_SPTA_ of (**a**) 155 mW/cm^2^ and (**b**) 478 mW/cm^2^. The minimum rectifier output voltage (V_rect_) required to operate the sensor was ∼1.36 V (**a**). The maximum V_rect_ that can be generated by the IC was ∼3 V limited by voltage limiting clamps at the rectifier input to prevent breakdown of the transistors (**b**).

**Supplementary Fig. 13.**
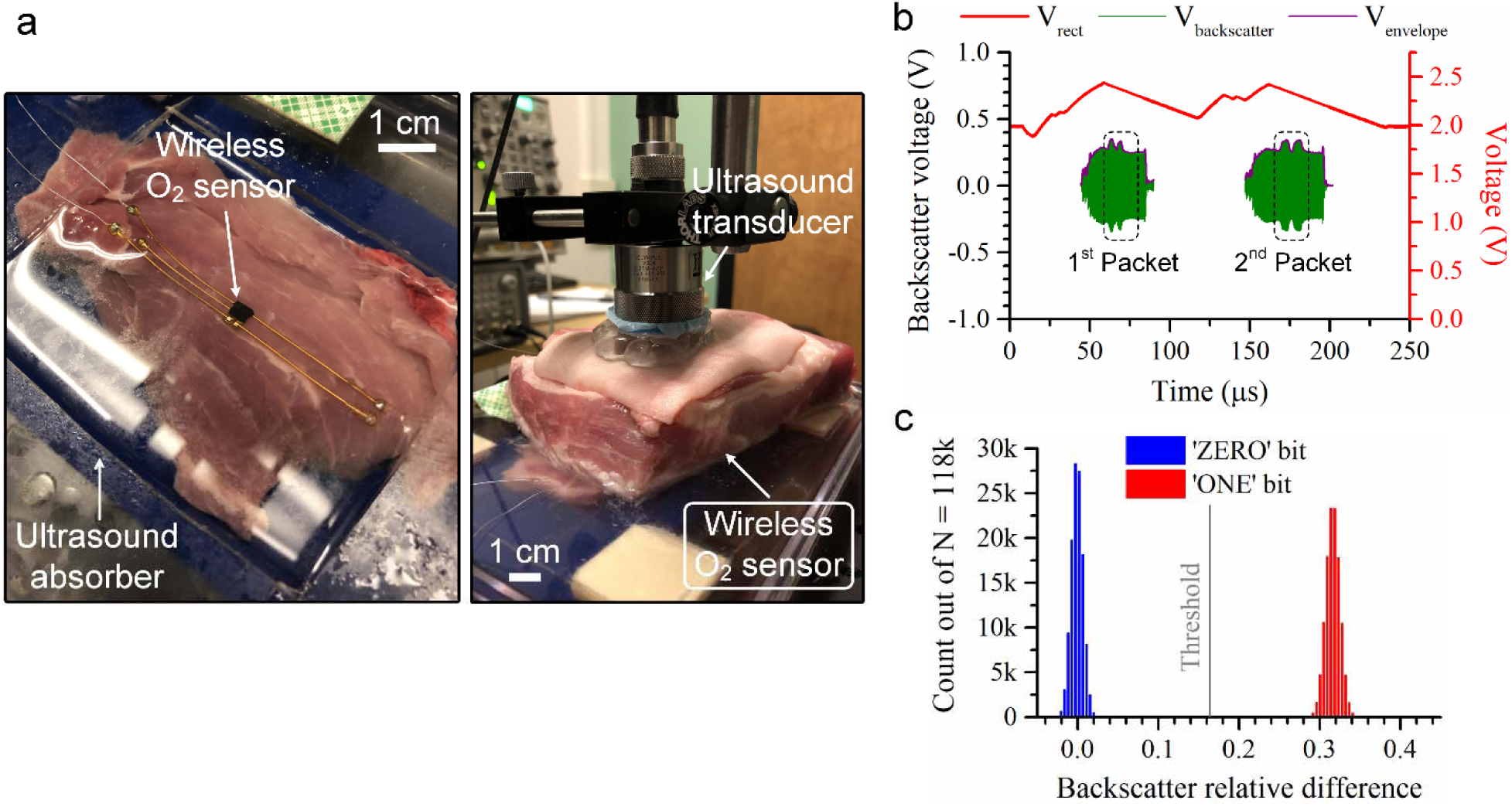
**a,** The wireless O_2_ sensor was at 5 cm depth through a fresh, ex vivo porcine tissue specimen, in which ultrasound waves with 660 mW/cm^2^ de-rated I_SPTA_, producing an acoustic power of ∼27.67 mW at the external transducer surface, propagated through approximately 2 mm ultrasound gel, 1.5 mm skin, 1 mm fat, and 45.5 mm muscle tissue. **b,** Sensor waveform and backscatter signal were captured in the wireless measurement of a single O_2_ sample. **c,** Backscatter relative difference for 118k O_2_ samples, showing ∼32% modulation depth. The system achieved a bit error rate (BER) of < 10^-5^ and a wireless link power transfer efficiency of ∼0.73%. The modulation depth measured ex vivo was lower than the modulation depth measured in vitro in distilled (DI) water (see Fig. 4d) due to a decrease in the ratio of the modulation amplitude (that is, the amplitude difference between the modulated and unmodulated backscatter signals) to the amplitude of the unmodulated backscatter signal. This ratio decrease can be attributed to the ultrasound reflections from internal tissue interfaces that interfered with the total US reflections from the sensor’s piezo and the part of the sensor surface at the face of the external transducer. Note that to make a fair comparison, the alignment between central axes of the piezo and the acoustic field was well-tuned by monitoring the rectifier voltage (V_rect_) amplitude in this measurement and the measurement in DI water.

**Supplementary Fig. 14.**
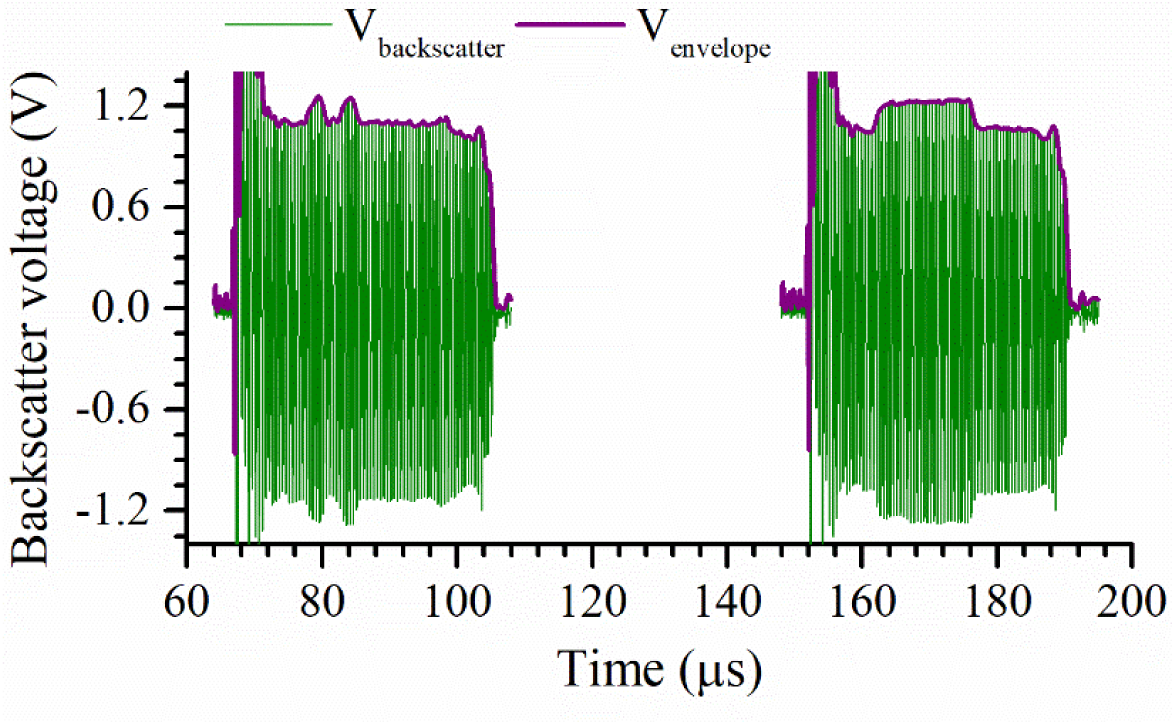
A backscatter signal from the wireless O_2_ sensor captured in an *in vivo* experiment, showing a high modulation depth of ∼15%.

**Supplementary Fig. 15.**
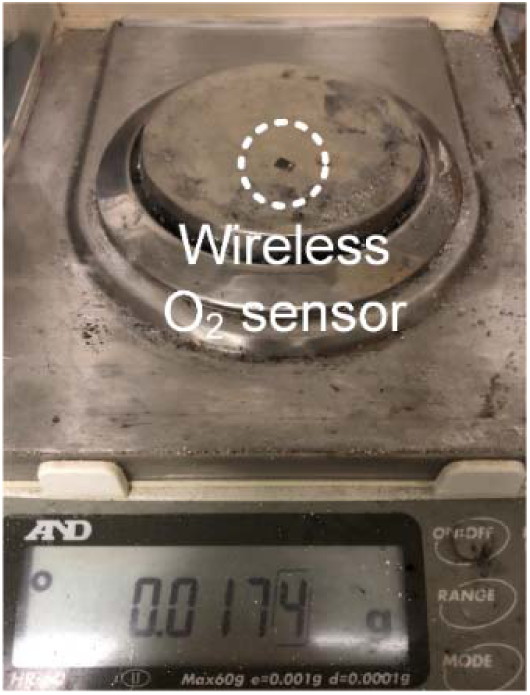
Photograph of a fully implantable, wireless, battery-free luminescence sensor on balance.

**Supplementary Fig. 16.**
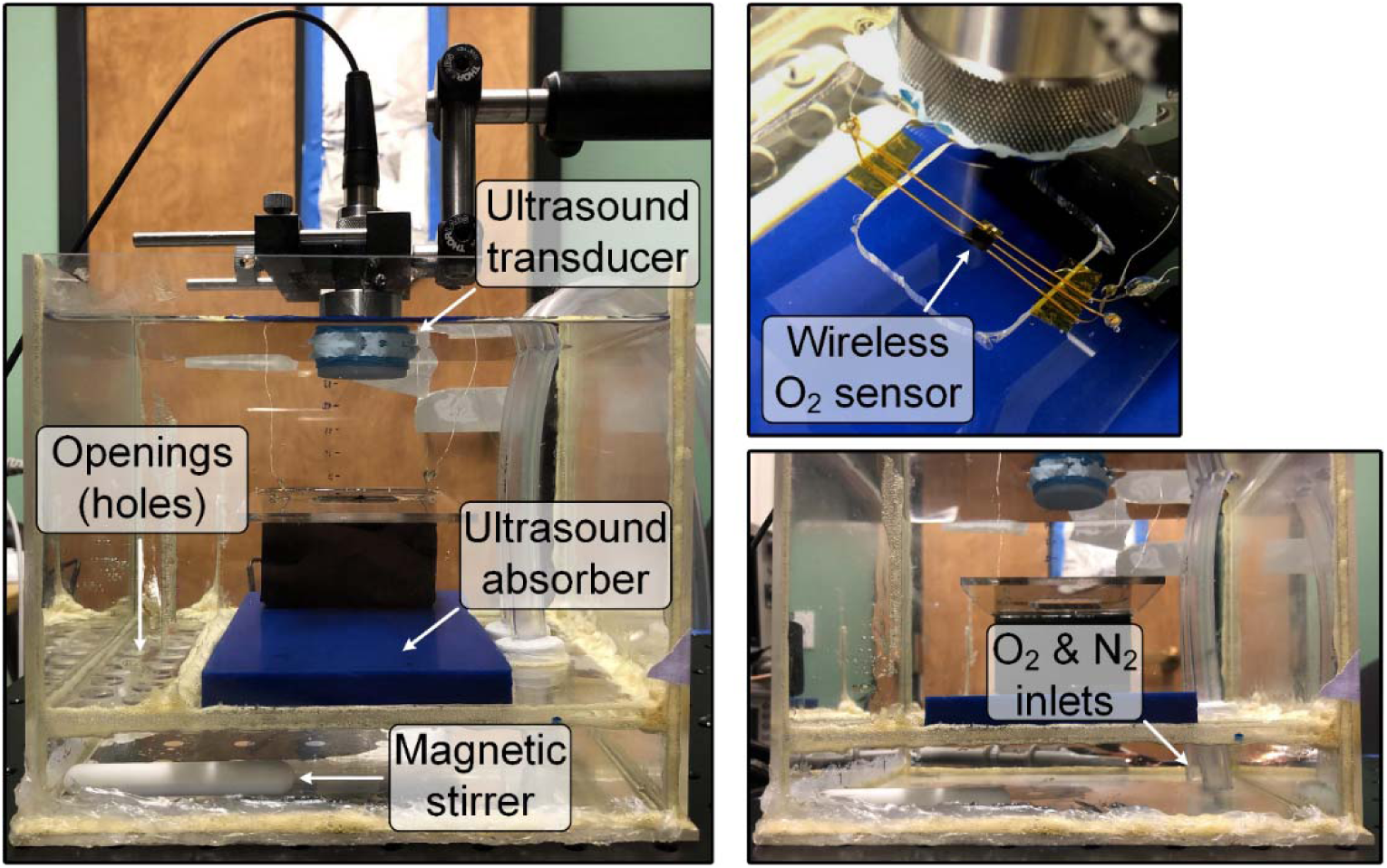
Photograph of the custom-built water tank used for in vitro characterization of the wireless system. The tank consisted of two separate compartments that are connected through openings (holes). A magnetic stirrer in the bottom compartment facilitated the mixing of oxygen (O_2_) and nitrogen (N_2_), purged through the pipes, with water. An ultrasound absorber placed to the bottom surface of the top compartment to reduce ultrasound reflections from the bottom surface during measurements.

**Supplementary Fig. 18.**
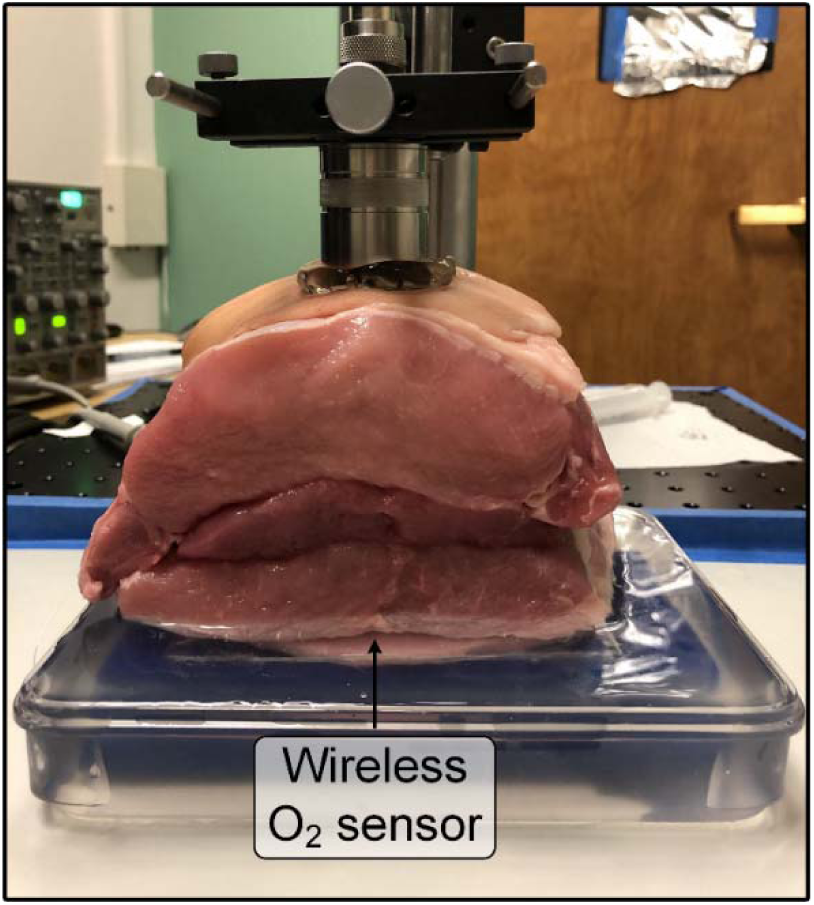
An alternate photograph for Fig. 6g taken from a different perspective.

**Supplementary Fig. 19.**
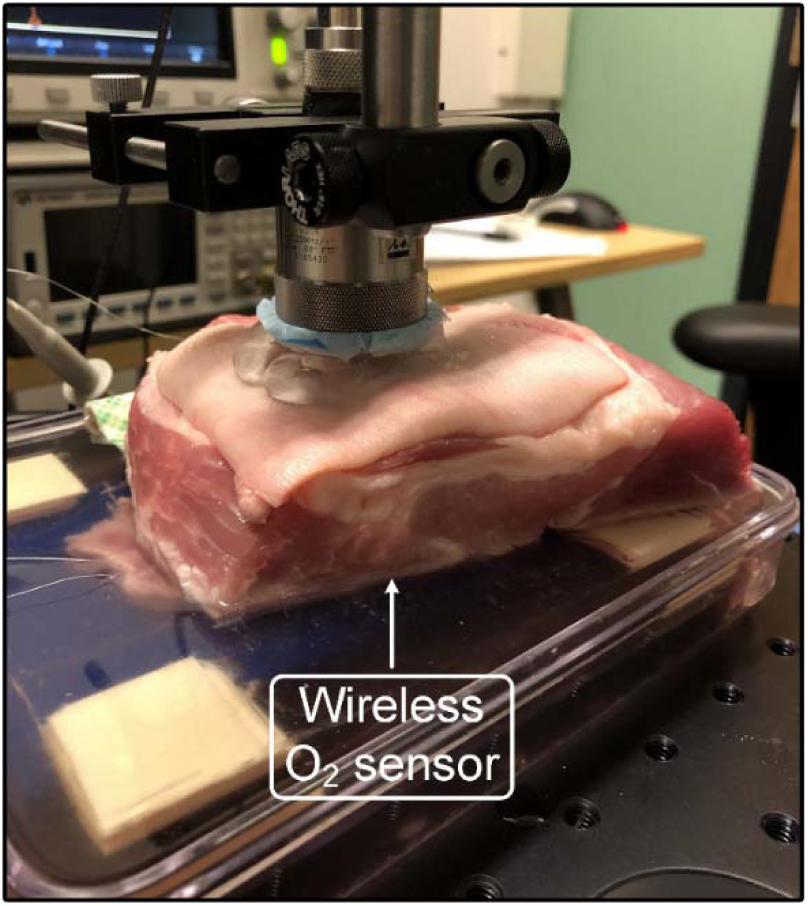
An alternate photograph for Supplementary Fig. 13a taken from a different perspective.

**Supplementary Fig. 20.**
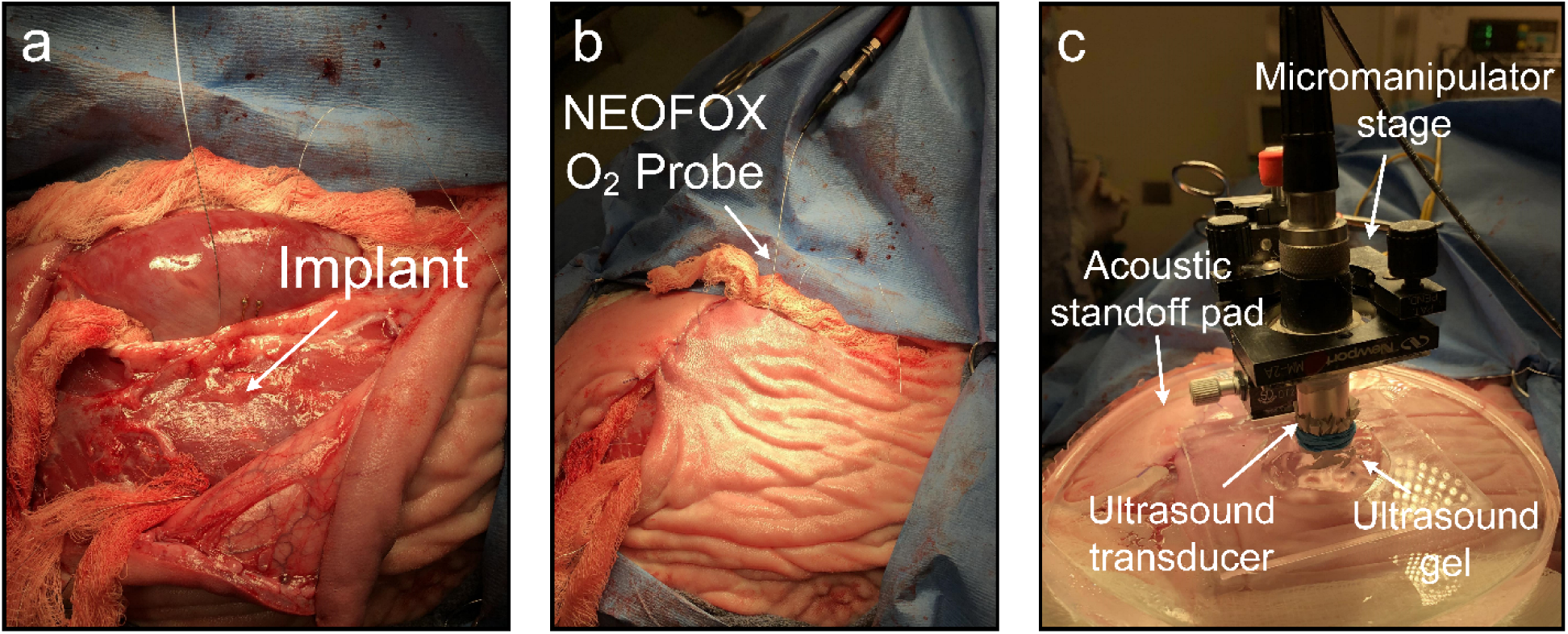
Photographs from animal B, showing the surgical placement of the implantable, wireless O_2_ sensor in a sheep model. See Fig. 5a-d caption for descriptions.

## References

1. Gibbons, R. D., Meltzer, D. & Duan, N. Waiting for Organ Transplantation. Science 287, 237 (2000).

2. Piardi, T. Vascular complications following liver transplantation: A literature review of advances in 2015. World J. Hepatol. 8, 36 (2016).

3. Mitchell, R. N. Graft Vascular Disease: Immune Response Meets the Vessel Wall. Annu. Rev. Pathol. Mech. Dis. 4, 19–47 (2009).

4. van den Brink, W. A. et al. Brain Oxygen Tension in Severe Head Injury. Neurosurgery 46, 868–878 (2000).

5. Gray, M. E. et al. In vivo validation of a miniaturized electrochemical oxygen sensor for measuring intestinal oxygen tension. Am. J. Physiol.-Gastrointest. Liver Physiol. 317, G242–G252 (2019).

6. Peterson, J. I., Fitzgerald, R. V. & Buckhold, D. K. Fiber-optic probe for in vivo measurement of oxygen partial pressure. Anal. Chem. 56, 62–67 (1984).

7. Lebedev, A. Y. et al. Dendritic Phosphorescent Probes for Oxygen Imaging in Biological Systems. ACS Appl. Mater. Interfaces 1, 1292–1304 (2009).

8. Lukina, M. et al. Interrogation of metabolic and oxygen states of tumors with fiber-based luminescence lifetime spectroscopy. Opt. Lett. 42, 731–734 (2017).

9. Marland, J. R. K. et al. Real-time measurement of tumour hypoxia using an implantable microfabricated oxygen sensor. https://osf.io/fhqd7 (2020) doi:10.31224/osf.io/fhqd7.

10. Nöth, U. et al. In vivo determination of tumor oxygenation during growth and in response to carbogen breathing using 15C5-loaded alginate capsules as fluorine-19 magnetic resonance imaging oxygen sensors. *Int*. J. Radiat. Oncol. 60, 909–919 (2004).

11. Liu, S. et al. Quantitative Tissue Oxygen Measurement in Multiple Organs Using 19F MRI in a Rat Model. Magn. Reson. Med. Off. J. Soc. Magn. Reson. Med. Soc. Magn. Reson. Med. 66, 1722–1730 (2011).

12. Elas, M. et al. Electron Paramagnetic Resonance Oxygen Images Correlate Spatially and Quantitatively with Oxylite Oxygen Measurements. Clin. Cancer Res. 12, 4209–4217 (2006).

13. Khan, N., Williams, B. B., Hou, H., Li, H. & Swartz, H. M. Repetitive Tissue pO2 Measurements by Electron Paramagnetic Resonance Oximetry: Current Status and Future Potential for Experimental and Clinical Studies. Antioxid. Redox Signal. 9, 1169–1182 (2007).

14. Gómez, H. et al. Use of non-invasive NIRS during a vascular occlusion test to assess dynamic tissue O2 saturation response. Intensive Care Med. 34, 1600 (2008).

15. Lynch, J. M. et al. Noninvasive Optical Quantification of Cerebral Venous Oxygen Saturation in Humans. Acad. Radiol. 21, 162–167 (2014).

16. Gladytz, T. et al. Near infrared spectroscopy system for quantitative monitoring of renal hemodynamics and oxygenation in rats. in Optical Tomography and Spectroscopy of Tissue XIII vol. 10874 108740H (International Society for Optics and Photonics, 2019).

17. Yamakawa, T. et al. Development of an Implantable Flexible Probe for Simultaneous Near-Infrared Spectroscopy and Electrocorticography. IEEE Trans. Biomed. Eng. 61, 388– 395 (2014).

18. Zhang, H. et al. Wireless, battery-free optoelectronic systems as subdermal implants for local tissue oximetry. Sci. Adv. 5, eaaw0873 (2019).

19. Ho, J. S. et al. Wireless power transfer to deep-tissue microimplants. Proc. Natl. Acad. Sci. 111, 7974–7979 (2014).

20. Seo, D. et al. Wireless Recording in the Peripheral Nervous System with Ultrasonic Neural Dust. Neuron 91, 529–539 (2016).

21. Piech, D. K. et al. A wireless millimetre-scale implantable neural stimulator with ultrasonically powered bidirectional communication. *Nat*. Biomed. Eng. 4, 207–222 (2020).

22. Wang, M. L. et al. Wireless data links for next-generation networked micro-implantables. in 2018 IEEE Custom Integrated Circuits Conference (CICC) 1–9 (2018). doi:10.1109/CICC.2018.8357096.

23. Thimot, J. & Shepard, K. L. Bioelectronic devices: Wirelessly powered implants. *Nat*. Biomed. Eng. 1, (2017).

24. O’Leary, B. & Vaezy, S. Marketing Clearance of Diagnostic Ultrasound Systems and Transducers. 64.

25. Ammi, A. Y. et al. Characterization of Ultrasound Propagation Through Ex-vivo Human Temporal Bone. Ultrasound Med. Biol. 34, 1578–1589 (2008).

26. Seo, D. et al. Ultrasonic beamforming system for interrogating multiple implantable sensors. in 2015 37th Annual International Conference of the IEEE Engineering in Medicine and Biology Society (EMBC) 2673–2676 (2015). doi:10.1109/EMBC.2015.7318942.

27. Maleki, T. et al. An Ultrasonically Powered Implantable Micro-Oxygen Generator (IMOG). IEEE Trans. Biomed. Eng. 58, 3104–3111 (2011).

28. Kim, A. et al. An Implantable Ultrasonically-Powered Micro-Light-Source (µLight) for Photodynamic Therapy. Sci. Rep. 9, (2019).

29. Shi, C., Costa, T., Elloian, J., Zhang, Y. & Shepard, K. L. A 0.065-mm3 Monolithically- Integrated Ultrasonic Wireless Sensing Mote for Real-Time Physiological Temperature Monitoring. IEEE Trans. Biomed. Circuits Syst. 14, 412–424 (2020).

30. Weber, M. J. et al. A Miniaturized Single-Transducer Implantable Pressure Sensor With Time-Multiplexed Ultrasonic Data and Power Links. IEEE J. Solid-State Circuits 53, 1089–1101 (2018).

31. Chodavarapu, V. P. et al. CMOS-Based Phase Fluorometric Oxygen Sensor System. IEEE Trans. Circuits Syst. Regul. Pap. 54, 111–118 (2007).

32. Yao, L., Khan, R., Chodavarapu, V. P., Tripathi, V. S. & Bright, F. V. Sensitivity-Enhanced CMOS Phase Luminometry System Using Xerogel-Based Sensors. IEEE Trans. Biomed. Circuits Syst. 3, 304–311 (2009).

33. McDonagh, C. et al. Phase fluorometric dissolved oxygen sensor. Sens. Actuators B Chem. 74, 124–130 (2001).

34. Jorge, P. A. S., Caldas, P., Rosa, C. C., Oliva, A. G. & Santos, J. L. Optical fiber probes for fluorescence based oxygen sensing. Sens. Actuators B Chem. 103, 290–299 (2004).

35. Yang, S. et al. Thermally resistant UV-curable epoxy–siloxane hybrid materials for light emitting diode (LED) encapsulation. J. Mater. Chem. 22, 8874–8880 (2012).

36. Wang, X. & S. Wolfbeis, O. Optical methods for sensing and imaging oxygen: materials, spectroscopies and applications. Chem. Soc. Rev. 43, 3666–3761 (2014).

37. Holland, R. Resonant Properties of Piezoelectric Ceramic Rectangular Parallelepipeds. J. Acoust. Soc. Am. 43, 988–997 (1968).

38. Goss, S. A., Frizzell, L. A. & Dunn, F. Ultrasonic absorption and attenuation in mammalian tissues. Ultrasound Med. Biol. 5, 181–186 (1979).

39. Chan, C.-M., Chan, M.-Y., Zhang, M., Lo, W. & Wong, K.-Y. The performance of oxygen sensing films with ruthenium-adsorbed fumed silica dispersed in silicone rubber. Analyst 124, 691–694 (1999).

40. Lu, X. & Winnik, M. A. Luminescence Quenching in Polymer/Filler Nanocomposite Films Used in Oxygen Sensors. Chem. Mater. 13, 3449–3463 (2001).

41. Lu, X., Manners, I. & Winnik, M. A. Polymer/Silica Composite Films as Luminescent Oxygen Sensors. Macromolecules 34, 1917–1927 (2001).

42. Li, X.-M., Ruan, F.-C. & Wong, K.-Y. Optical characteristics of a ruthenium(II) complex immobilized in a silicone rubber film for oxygen measurement. Analyst 118, 289–292 (1993).

43. Pfeiffer, S. A. & Nagl, S. Microfluidic platforms employing integrated fluorescent or luminescent chemical sensors: a review of methods, scope and applications. Methods Appl. Fluoresc. 3, 034003 (2015).

44. Morris, K. J., Roach, M. S., Xu, W., Demas, J. N. & DeGraff, B. A. Luminescence Lifetime Standards for the Nanosecond to Microsecond Range and Oxygen Quenching of Ruthenium(II) Complexes. Anal. Chem. 79, 9310–9314 (2007).

45. Hartmann, P., Leiner, M. J. P. & Kohlbacher, P. Photobleaching of a ruthenium complex in polymers used for oxygen optodes and its inhibition by singlet oxygen quenchers. Sens. Actuators B Chem. 51, 196–202 (1998).

46. Sonmezoglu, S. & Maharbiz, M. M. 34.4 A 4.5mm3 Deep-Tissue Ultrasonic Implantable Luminescence Oxygen Sensor. in 2020 IEEE International Solid- State Circuits Conference - (ISSCC) 454–456 (2020). doi:10.1109/ISSCC19947.2020.9062946.

47. Guo, J. & Sonkusale, S. A 65 nm CMOS Digital Phase Imager for Time-Resolved Fluorescence Imaging. IEEE J. Solid-State Circuits 47, 1731–1742 (2012).

48. Zhang, X. & Apsel, A. B. A Low-Power, Process-and- Temperature- Compensated Ring Oscillator With Addition-Based Current Source. IEEE Trans. Circuits Syst. Regul. Pap. 58, 868–878 (2011).

49. Richards, P. L. Bolometers for infrared and millimeter waves. J. Appl. Phys. 76, 1–24 (1994).

50. Ozeri, S. & Shmilovitz, D. Simultaneous backward data transmission and power harvesting in an ultrasonic transcutaneous energy transfer link employing acoustically dependent electric impedance modulation. Ultrasonics 54, 1929–1937 (2014).

51. Ghanbari, M. M. et al. A Sub-mm3 Ultrasonic Free-Floating Implant for Multi-Mote Neural Recording. IEEE J. Solid-State Circuits 54, 3017–3030 (2019).

52. Kino, G. S. Acoustic Waves: Devices, Imaging, and Analog Signal Processing. (Prentice Hall, 1987).

53. El-Sheimy, N., Hou, H. & Niu, X. Analysis and Modeling of Inertial Sensors Using Allan Variance. IEEE Trans. Instrum. Meas. 57, 140–149 (2008).

54. Carraway, E. R., Demas, J. N., DeGraff, B. A. & Bacon, J. R. Photophysics and photochemistry of oxygen sensors based on luminescent transition-metal complexes. Anal. Chem. 63, 337–342 (1991).

55. Adkins, J. N. et al. Toward a Human Blood Serum Proteome: Analysis By Multidimensional Separation Coupled With Mass Spectrometry. Mol. Cell. Proteomics 1, 947–955 (2002).

56. Ratner, B. D. Replacing and Renewing: Synthetic Materials, Biomimetics, and Tissue Engineering in Implant Dentistry. J. Dent. Educ. 65, 1340–1347 (2001).

57. Blaszykowski, C., Sheikh, S. & Thompson, M. Surface chemistry to minimize fouling from blood-based fluids. Chem. Soc. Rev. 41, 5599–5612 (2012).

58. Rudolph, A. M. Circulatory changes during gestational development of the sheep and human fetus. Pediatr. Res. 84, 348–351 (2018).

59. Samson, N., Fortin-Pellerin, E. & Praud, J.-P. The contribution of ovine models to perinatal respiratory physiology. Front. Biosci. Landmark Ed. 23, 1195–1219 (2018).

60. Drury, P. P., Gunn, E. R., Bennet, L. & Gunn, A. J. Mechanisms of Hypothermic Neuroprotection. Clin. Perinatol. 41, 161–175 (2014).

61. Yates, D. T. et al. Myoblasts from intrauterine growth-restricted sheep fetuses exhibit intrinsic deficiencies in proliferation that contribute to smaller semitendinosus myofibres. J. Physiol. 592, 3113–3125 (2014).

62. Grant, S. A., Bettencourt, K., Krulevitch, P., Hamilton, J. & Glass, R. In vitro and in vivo measurements of fiber optic and electrochemical sensors to monitor brain tissue pH. Sens. Actuators B Chem. 72, 174–179 (2001).

63. Borisov, S. M., Krause, C., Arain, S. & Wolfbeis, O. S. Composite Material for Simultaneous and Contactless Luminescent Sensing and Imaging of Oxygen and Carbon Dioxide. Adv. Mater. 18, 1511–1516 (2006).

64. DeHennis, A., Getzlaff, S., Grice, D. & Mailand, M. An NFC-Enabled CMOS IC for a Wireless Fully Implantable Glucose Sensor. IEEE J. Biomed. Health Inform. 20, 18–28 (2016).

65. Wang, X., S. Wolfbeis, O. & J. Meier, R. Luminescent probes and sensors for temperature. Chem. Soc. Rev. 42, 7834–7869 (2013).

66. Chen, X. et al. Recent progress in the development of fluorescent, luminescent and colorimetric probes for detection of reactive oxygen and nitrogen species. Chem. Soc. Rev. 45, 2976–3016 (2016).

67. National Data - OPTN. https://optn.transplant.hrsa.gov/data/view-data-reports/national-data/.

68. Norton, P. T., DeAngelis, G. A., Ogur, T., Saad, W. E. & Hagspiel, K. D. Noninvasive Vascular Imaging in Abdominal Solid Organ Transplantation. Am. J. Roentgenol. 201, W544–W553 (2013).

69. Boraschi, P., Pina, M. C. D. & Donati, F. Graft complications following orthotopic liver transplantation: Role of non-invasive cross-sectional imaging techniques. Eur. J. Radiol. 85, 1271–1283 (2016).

70. Hickman, P. E., Potter, J. M. & Pesce, A. J. Clinical chemistry and post-liver-transplant monitoring. Clin. Chem. 43, 1546–1554 (1997).

71. Khaja, M. S., Matsumoto, A. H. & Saad, W. E. Complications of Transplantation. Part 1: Renal Transplants. Cardiovasc. Intervent. Radiol. 37, 1137–1148 (2014).

72. Wunsch, H., Angus, D. C., Harrison, D. A., Linde-Zwirble, W. T. & Rowan, K. M. Comparison of Medical Admissions to Intensive Care Units in the United States and United Kingdom. Am. J. Respir. Crit. Care Med. 183, 1666–1673 (2011).

73. Barrett, M. L., Smith, M. W., Elixhauser, A., Honigman, L. S. & Pines, J. M. Utilization of Intensive Care Services, 2011: Statistical Brief #185. in Healthcare Cost and Utilization Project (HCUP) Statistical Briefs (Agency for Healthcare Research and Quality (US), 2006).

74. Kara, A., Akin, S. & Ince, C. Monitoring microcirculation in critical illness. Curr. Opin. Crit. Care 22, (2016).

75. Dyson, A. & Singer, M. Tissue oxygen tension monitoring: will it fill the void? Curr. Opin. Crit. Care 17, (2011).

76. Brandstrup, B. et al. Effects of Intravenous Fluid Restriction on Postoperative Complications: Comparison of Two Perioperative Fluid Regimens: A Randomized Assessor-Blinded Multicenter Trial. Ann. Surg. 238, (2003).

77. Dünser, M. W. et al. Association of arterial blood pressure and vasopressor load with septic shock mortality: a post hoc analysis of a multicenter trial. Crit. Care 13, R181 (2009).

78. Ince, C. Hemodynamic coherence and the rationale for monitoring the microcirculation. Crit. Care 19, S8 (2015).

79. Wang, M. L., Chang, T. C. & Arbabian, A. Ultrasonic Implant Localization for Wireless Power Transfer: Active Uplink and Harmonic Backscatter. in 2019 IEEE International Ultrasonics Symposium (IUS) 818–821 (2019). doi:10.1109/ULTSYM.2019.8926006.

80. Koch, T. et al. Ultrasound velocity and attenuation of porcine soft tissues with respect to structure and composition: I. Muscle. Meat Sci. 88, 51–58 (2011).

81. Wang, M. L. et al. Closed-loop ultrasonic power and communication with multiple miniaturized active implantable medical devices. in 2017 IEEE International Ultrasonics Symposium (IUS) 1–4 (2017). doi:10.1109/ULTSYM.2017.8092116.

82. Shen, K. & Maharbiz, M. M. Ceramic Packaging in Neural Implants. bioRxiv 2020.06.26.174144 (2020) doi:10.1101/2020.06.26.174144.

83. Shen, K. & Maharbiz, M. M. Design of Ceramic Packages for Ultrasonically Coupled Implantable Medical Devices. IEEE Trans. Biomed. Eng. 67, 2230–2240 (2020).

84. Traeger, R. Nonhermeticity of Polymeric Lid Sealants. IEEE Trans. Parts Hybrids Packag. 13, 147–152 (1977).

85. Song, E., Li, J., Won, S. M., Bai, W. & Rogers, J. A. Materials for flexible bioelectronic systems as chronic neural interfaces. Nat. Mater. 19, 590–603 (2020).

86. Hassler, C., Boretius, T. & Stieglitz, T. Polymers for neural implants. J. Polym. Sci. Part B Polym. Phys. 49, 18–33 (2011).

87. Chang, T. C. et al. Design of Tunable Ultrasonic Receivers for Efficient Powering of Implantable Medical Devices With Reconfigurable Power Loads. IEEE Trans. Ultrason. Ferroelectr. Freq. Control 63, 1554–1562 (2016).

88. Hughes, S. W. Archimedes revisited: a faster, better, cheaper method of accurately measuring the volume of small objects. Phys. Educ. 40, 468–474 (2005).

89. Sato, Y., Mizutani, K., Wakatsuki, N. & Nakamura, T. Design for an Aspherical Acoustic Fresnel Lens with Phase Continuity. Jpn. J. Appl. Phys. 47, 4354–4359 (2008).

90. Sanchis, L., Yánez, A., Galindo, P. L., Pizarro, J. & Pastor, J. M. Three-dimensional acoustic lenses with axial symmetry. Appl. Phys. Lett. 97, 054103 (2010).

